# Mechanistic Insight into AP-Endonuclease 1 Cleavage of Abasic Sites at Stalled Replication Forks

**DOI:** 10.1101/2022.08.22.504733

**Authors:** Nicole M. Hoitsma, Jessica Norris, Thu H. Khoang, Vikas Kaushik, Edwin Antony, Mark Hedglin, Bret D. Freudenthal

## Abstract

Many types of DNA damage stall replication fork progression, including abasic sites. AP-Endonuclease 1 (APE1) has been shown to cleave abasic sites in ssDNA substrates. Importantly, APE1 cleavage of ssDNA at a replication fork has significant biological implications by generating double strand breaks that could collapse the replication fork. Despite this, the molecular basis and efficiency of APE1 processing abasic sites at a replication fork remains elusive. Here, we investigate APE1 cleavage of several abasic substrates that mimic potential APE1 interactions at replication forks. We determine that APE1 has robust activity on these substrates, similar to dsDNA, and report rapid rates for cleavage and product release. X-ray crystal structures visualize the APE1 active site, highlighting that a similar mechanism is used to process ssDNA substrates as canonical APE1 activity on dsDNA. However, mutational analysis reveals R177 to be uniquely critical for the APE1 ssDNA cleavage mechanism. Additionally, we investigate the interplay between APE1 and Replication Protein A (RPA), the major ssDNA-binding protein at replication forks, revealing that APE1 can cleave an abasic site while RPA is still bound to the DNA substrate. Together, this work provides molecular level insights into abasic ssDNA processing by APE1, including the presence of RPA.

## 2. Introduction

Abasic sites are one of the most frequent types of DNA damage, with an estimated occurrence of ∼10,000-20,000 per human cell per day. These non-coding lesions are generated through spontaneous depurination or enzymatic processing by DNA glycosylases, which remove the nitrogenous base and leave behind a baseless sugar moiety^1–4^. Abasic sites preferentially form at sites of DNA replication, as ssDNA is more vulnerable to chemical attack and spontaneous base loss^3, 5^. Furthermore, APOBEC enzymes preferentially deaminate cytosines in ssDNA and, with subsequent processing by uracil N-glycosylase (UNG), contribute to abasic site accumulation in ssDNA at replication forks^3, 6–11^. This poses a threat since abasic sites act as a block to numerous essential cellular processes, including transcription and replication. When replicative polymerases encounter abasic sites, they pause at the lesion, ultimately stalling the replication fork at a 3′ dsDNA-ssDNA primer-template junction (PTJ)^12–16^. However, how abasic sites at stalled replication forks are targeted to distinct bypass/repair pathways remains unknown. As highlighted in **Figure 1**, processing can occur via a number of pathways: (1) the abasic site can be cleaved by the nuclease Apurinic/apyrimidinic endonuclease 1 (APE1) generating a double strand DNA break, (2) the abasic site can be covalently attached to the HMCES protein generating a DNA protein crosslink^11, 17–20^, or (3) the abasic site can be bypassed by a translesion DNA polymerase generating mutations due to the lack of coding potential^18, 21–22^.

**Figure 1.**
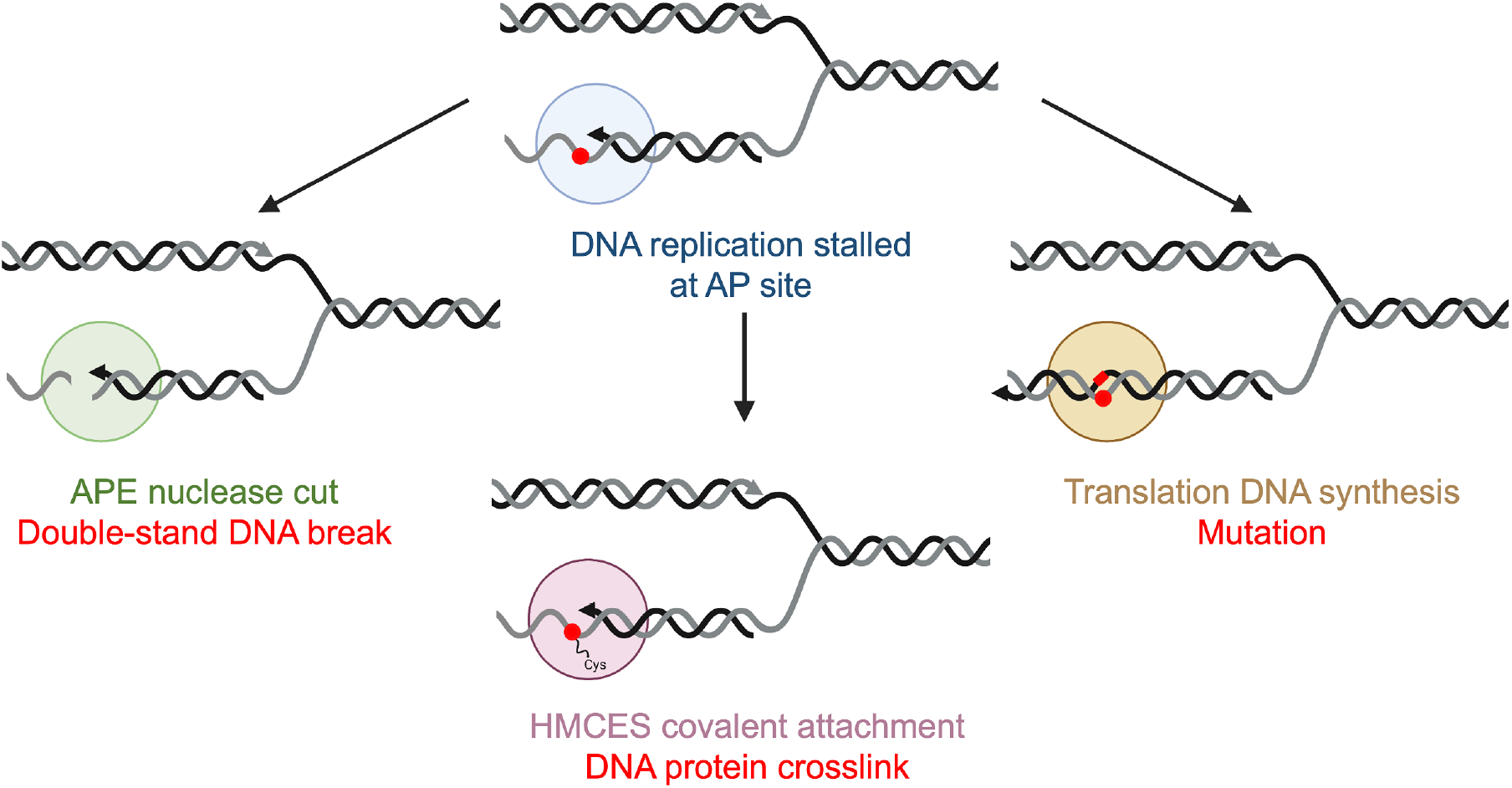
Pathways for AP site repair/bypass at a stalled replication fork. When the replicative DNA polymerase (blue, top) encounters a AP site (red sphere), it will stall the replication fork and processing can occur via a number of pathways: (1) the abasic site can be cleaved by the nuclease APE1 (green, left) generating a double strand DNA break, (2) the abasic site can be covalently attached to the HMCES protein (pink, bottom) generating a DNA protein crosslink, or (3) the abasic site can be bypassed by a translation DNA polymerase (yellow, right) generating mutations due to the lack of coding potential. Figure created with BioRender.com.

As an essential enzyme in the base excision repair pathway, APE1 has a well characterized cleavage activity processing abasic sites in double stranded DNA (dsDNA)^23–24^. However, APE1 also cleaves abasic sites in a variety of other biologically relevant substrates, including single stranded DNA (ssDNA), thus any abasic site near the replication fork is a potential cleavage site for APE1^25–26^. Despite the biological implications of APE1 cleavage at the replication fork, the molecular basis for APE1 recognizing and processing abasic sites in substrates mimicking the replication fork remains unknown. This is particularly important given the abundant concentration of cellular APE1 and data showing physical association of APE1 with components of the replication machinery^27–29^. Furthermore, evidence that there is an APE1-dependent degradation of stalled replication forks in response to DNA damage demonstrates the significance of APE1 in replication fork stability^30^.

APE1 cleavage of ssDNA has significant biological implications at a stalled replication fork because it generates a double stranded break, which could collapse the replication fork. In this way, it is likely that there are protections against this APE1 activity in cells, such as inhibition by single-stranded DNA binding proteins that quickly and tightly bind to ssDNA within the cell. In particular, replication protein A (RPA), which is the major ssDNA-binding protein in cells, binds to ssDNA with very high affinity. RPA has been identified as a negative regulator of APE1 ssDNA cleavage activity suggesting that RPA may provide a mechanism for regulating APE1 activity in cells^26^. While RPA does not interact directly with human APE1, RPA has high affinity for ssDNA, even with an abasic site, leading to the hypothesis that RPA may inhibit APE1 ssDNA cleavage by coating the ssDNA and preventing APE1:ssDNA complex formation^26^. However, the molecular basis and efficiency for APE1 processing abasic sites in substrates mimicking a replication fork, as well as the influence of RPA, remain significant outstanding questions.

Here, we study APE1 cleavage activity at abasic DNA substrates that mimic potential APE1 interactions at a stalled replication fork using a combination of kinetic and structural approaches. We determine that APE1 has robust activity on these replication fork mimic substrates, reporting pre-steady-state kinetic rates for both cleavage and product release. X-ray crystallography reveals molecular details of the APE1 active site during catalysis, including active site contacts and residues that mediate cleavage of abasic sites in ssDNA, highlighting that APE1 uses a similar mechanism to process ssDNA and dsDNA substrates. Through mutational and kinetic analysis, we determine residue R177 to be critical for the APE1 ssDNA cleavage mechanism. Furthermore, we investigated the interplay between APE1 and RPA on these stalled replication fork substrates. Data reveals that APE1 is able to catalyze cleavage of the phosphodiester backbone with RPA bound to the DNA substrate. Together these data provide insight into the complex role of APE1 at the stalled replication forks, including its activity in the presence of RPA.

## 3. Materials and Methods

### 3.1 DNA Sequences

DNA oligonucleotides were synthesized by Integrated DNA Technologies (Coralville, IA) and annealed as described below. An abasic site analog, tetrahydrofuran (THF), was utilized for stability and is denoted in the following DNA sequences as an underlined <u>X</u>. Primer template junction (PTJ) substrates were generated by annealing the following sequences: damaged strand with a 5′ fluorescein label(indicated by the asterisk) 5′-*TGGATGATGACTCTTCTGCGTTCGCTGATGCGC<u>X</u>CGACGGTAGTTAAGTGTT GAG-3′, non-damaged strand, 5′-CTCAACACTTAACTACCGTCG-3′ (for PTJ) and 5′-CTCAACACTTAACTA-3′ (for Recessed PTJ). The ssDNA substrate sequence used for kinetics and binding assays was: 5′-*GCT-GAT-GCG-CXC-GAC-GGA-TCC-3′. For crystallization of the APE1:ssDNA complex the following ssDNA sequence was used: 5′-ATCCGAFCGATGC-3′. After resuspension, the single stranded oligonucleotide concentrations were determined by absorbance at 260 nm. Annealing reactions were completed by mixing equimolar amounts of the complimentary oligonucleotides, heating to 95°C for 5 minutes, and then allowing the reaction to cool at a rate of 1°C min^−1^ to 4°C.

For FRET experiments, oligonucleotides were purified on denaturing polyacrylamide gels. The concentrations of unlabeled DNAs were determined from the absorbance at 260 nm using the calculated extinction coefficients. The concentrations of Cy3-labeled DNAs were determined from the extinction coefficient at 550 nm for Cy3 (ε_550_ = 136,000 M^−1^cm^−1^). For annealing two single strand DNAs (as depicted in **Figure S1**), the primer and corresponding complementary template strands were mixed in equimolar amounts in 1X Annealing Buffer (10 mM TrisHCl, pH 8.0, 100 mM NaCl, 1 mM EDTA), heated to 95 °C for 5 minutes, and allowed to slowly cool to room temperature.

### 3.2 Protein expression and purification

Several different APE1 mutants were used in this study: Full length wild-type APE1 and full length APE1 R177A for activity assays, full length catalytically dead APE1 D210N/E96Q (APE1_Dead_) for binding and FRET assays, and truncated (lacking the N-terminal 42 amino acids) APE1 C138A (ΔAPE1) for crystallography. All of these APE1 proteins were expressed and purified as previously described^31^. Mutagenesis was done via Quick-change II site-directed mutagenesis (Agilent) and mutations were confirmed via Sanger sequencing. Plasmids, pet28a codon optimized, were overexpressed in BL21(DE3) plysS *E. coli* cells (Invitrogen). Expression was induced with isopropyl β-D-1-thiogalactopyranoside before cells were harvested and lysed via sonication. APE1 protein was purified from the cell lysate using column chromatography. The resins for this purification include heparin, cation exchange, and gel filtration resins on an ATKA-Pure FPLC. Purified protein was monitored throughout the purification process via SDS-PAGE. After column chromatography, pure fractions were pooled, and final concentration was determined via absorbance at 280 nm. Concentrated APE1 protein was stored at −80 °C in a buffer of 50 mM HEPES (pH 7.4) and 150 mM NaCl. Cy5-labeled human RPA was generated using non-canonical amino acids as described^32–33^.

### 3.3 APE1 activity assays

Reactions were initiated by mixing APE1 enzyme with DNA substrate in reaction buffer containing a final concentration of 50 mM HEPES, pH 7.5, 100 mM KCl, 5 mM MgCl2 and 0.1 mg/ml bovine serum albumin (BSA) at 37°C. Different APE1 and DNA final concentrations were used dependent upon the kinetic regime for each assay. For multiple turnover kinetics, DNA is in excess of APE1 at final concentrations of 100 and 30 nM, respectively. For single turnover kinetics, APE1 is in excess of DNA at final concentrations of 500 and 50 nM, respectively. For qualitative product formation assays, final concentrations were 5 nM APE1 and 500 nM DNA. After mixing, the reactions were allowed to progress for a pre-determined amount of time before being quenched with EDTA. Multiple and single turnover pre-steady-state kinetics were completed at very rapid timepoints, and thus were carried out using a rapid quench flow system (KinTek) which allows rapid mixing as fast as 0.002 sec. Temperature is controlled with a circulating water bath in the instrument allowing experiments to be carried out at 37° C. Longer time points used for the product formation assay (1 and 10 minutes) allowed reactions to be completed in a benchtop heat block at 37° C. Quenched reactions were mixed with loading dye (100 mM EDTA, 80% deionized formamide, 0.25 mg/ml bromophenol blue and 0.25 mg/ml xylene cyanol) and incubated at 95° C for 6 min. Reactions products were separated via denaturing polyacrylamide gel electrophoresis. DNA oligonucleotides used in this assay (described above) contain a 5′ fluorescein label (6-FAM) allowing substrate and product bands to be visualized with a Typhoon imager in fluorescence mode. Analysis of the imaged gel is completed by quantifying the bands using ImageQuant software, plotted and fit using Prism. To quantitatively determine rate values, multiple turnover time courses were fit to the following equation: Product = A(1−e^−*k*^_obs_^t^) + v_ss_t, where A represents the amplitude of the rising exponential which corresponds to the fraction of actively bound enzyme, and *k*_obs_ is the first order rate constant. The steady-state rate constant (*k*_ss_) is the steady-state velocity (v_ss_)/A. Single turnover experiments for wild-type APE1 are fit to single exponential equation: Product = A(1−e^−*k*^ ^t^). Single turnover experiments for R177A mutant are fit to double exponential equation: Product = A(1−e^−*k*^_1_^t^) + B(1−e^−*k*^_2_^t^), as seen previously^34^. Each time point in the curves for both single and multiple turnover kinetics represents an average of three independent experiments (technical replicates) ± standard error as determined using Prism analysis software.

### 3.4 Electrophoretic Mobility Shift Assay

Full length catalytically dead APE1 D210N/E96Q (APE1_Dead_) was utilized in binding experiments to prevent cleavage, which would convolute binding analysis. In these experiments, annealed or single stranded DNA substrates were mixed with varying amount of APE1 in reaction buffer containing a final concentration of 50 mM Tris (pH 8), 1 mM EDTA, 0.2 mg/ml BSA, 1 mM DTT, and 5% v/v sucrose (for purposes of gel loading). Samples are equilibrated at room temperature for 20 minutes before separation on a 10% native 59:1 polyacrylamide gel run at 120 V with 0.2X TBE running buffer. DNA oligonucleotides used in this assay (<u>described above</u>) contain a 5’ fluorescein label (6-FAM) which allows free and bound DNA bands to be visualized using a Typhoon imager in fluorescence mode. Analysis is completed by quantifying the free DNA bands using ImageQuant software, plotted and fit using Prism to Equation 1:

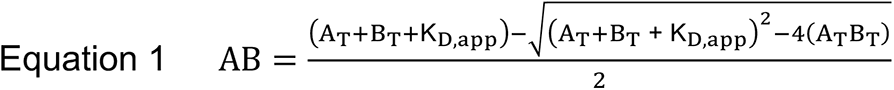

where A_T_ and B_T_ represent the total concentration of APE1 and DNA, respectively, and AB is the concentration of APE1:DNA complex used to determine apparent affinity (*K*_D_,_app_). Each data point represents an average of three independent experiments (technical replicates) ± standard error as determined using Prism software.

### 3.5 X-ray Crystallography

For crystallization of APE1:ssDNA product complex, the ssDNA oligo was mixed with ΔAPE1 C138A at a final concentration of 0.56 nM DNA and 10-12 mg/ml APE1. The truncation of N-terminal 42 amino acids and C138A mutation are commonly utilized to aid in APE1 crystallization^35–36^. As this is a catalytically competent APE1 enzyme, product formation likely occurs during a 30-minute incubation period at room temperature, prior to setting up crystallization trays. APE1:ssDNA product complex crystals were generated via sitting drop vapor diffusion using 2 ul protein/DNA mix combined with 2 ul reservoir solution (0.05M KCl; 0.05M sodium cacodylate pH 6; 10% PEG 8,000; 5mM spermine; 5mM L-Argininamide dihydrochloride). Resultant crystals were transferred to a cryoprotectant solution containing reservoir solution supplemented with 20% ethylene glycol, flash frozen, and subjected to X-ray diffraction. The APE1:ssDNA product structure was collected at 100 K on a Rigaku MicroMax-007 HF rotating anode diffractometer system at a wavelength of 1.54 Å. This system utilizes a Dectris Pilatus3R 200K-A detector and HKL3000R software was used for processing and scaling the data after collection. Initial models were determined using molecular replacement in PHENIX with a modified version of a previously determined APE1:DNA complex structure (PDB 5DFF). Refinement and model building were done with PHENIX and Coot, respectively, and figures were made using PyMol (Schrödinger LLC)^37–38^.

### 3.6 Ensemble FRET Measurements

All experiments were performed at room temperature (23 ± 2 °C) and, unless indicated otherwise, in 1X Mg^2+^/Ca^2+^ buffer (20 mM HEPES, pH 7.5, 150 mM KCl, 5 mM MgCl_2_, 5 mM CaCl_2_) supplemented with 1 mM DTT and the ionic strength was adjusted to physiological (200 mM) by the addition of appropriate amounts of KCl. All experiments were performed in a Horiba Scientific Duetta-Bio fluorescence/absorbance spectrometer. Solutions are excited at 514 nm and the fluorescence emission intensities (*I*) are simultaneously monitored at 563 nm (*I*_563_, Cy3 FRET donor fluorescence emission maximum) and 665 nm (*I*_665_, Cy5 FRET acceptor fluorescence emission maximum) over time, recording *I* every 0.17 s. For each time point, E_FRET_ is calculated where 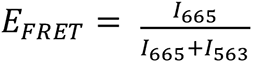. For all FRET experiments, excitation and emission slit widths are 20 nm. For RPA/APE1_Dead_ titration exchange experiments, a solution containing a 3′ Cy3-labeled PTJ DNA (**Figure S1**) is pre-saturated with Cy5-RPA and the resultant mixture is then titrated with increasing amounts of APE1^Dead^ and the E_FRET_ values at each protein amount is calculated as follows. At each protein amount, E_FRET_ values are recorded every 0.17 s until the FRET maintains a constant value for at least 1 minute and the data points within this stable region are averaged to obtain the final E_FRET_ value. To determine the predicted E_FRET_ value for Cy5-RPA remaining completely disengaged from the Cy3-PTJ DNA, the fluorescence emission intensities (*I*_665_ and *I*_563_) for Cy5-RPA alone and Cy3-PTJ DNA alone are each recorded over time, and E_FRET_ is calculated for each time point as follows; 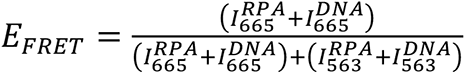. For RPA/DNA titration exchange experiments, a solution containing a Cy3-labeled Rec-PTJ DNA (with an abasic site 7 nt downstream of a PTJ, **Figure S1**) is pre-saturated with Cy5-RPA and the result mixture is then titrated with increasing amounts of an unlabeled DNA (either PTJ or Rec-PTJ DNA) and the E_FRET_ values at each DNA amount is calculated as described above. The predicted E_FRET_ value for Cy5-RPA remaining completely disengaged from the Cy3-PTJ DNA is determined as described above. Data for each titration is normalized to a range defined by the E_FRET_ observed in the absence of unlabeled competitor DNA (i.e., Y-Max) and the predicted E_FRET_ value for Cy5-RPA remaining completely disengaged from the Cy3-PTJ DNA (i.e., Y-Min). Each data point (E_FRET_ values) represents the mean ± S.E.M. of at least three independent measurements.

### 3.7 APE1 activity in the presence of RPA

These experimental conditions were designed to mimic the conditions of the FRET assay. All experiments were performed at room temperature in 1X Mg^2+^/Ca^2+^ buffer (20 mM HEPES, pH 7.5, 150 mM KCl, 5 mM MgCl_2_, 5 mM CaCl_2_) supplemented with 1 mM DTT and the ionic strength was adjusted to physiological (200 mM) by the addition of appropriate amounts of KCl. A solution containing the PTJ or Rec-PTJ DNA is pre-saturated with RPA protein. Reactions were initiated by mixing RPA-saturated DNA substrate with APE1 at varying concentrations (as indicated in the figure legends). After mixing, the reactions were allowed to progress for a pre-determined amount of time before being quenched with EDTA. Quenched reactions were mixed with loading dye (100 mM EDTA, 80% deionized formamide, 0.25 mg/ml bromophenol blue and 0.25 mg/ml xylene cyanol) and incubated at 95° C for 6 min. Reactions products were separated via denaturing polyacrylamide gel electrophoresis. DNA oligonucleotides used in this assay were the same as those used for kinetic assays and thus contain a 5′ fluorescein label (6-FAM) allowing substrate and product bands to be visualized with a Typhoon imager in fluorescence mode. Analysis of the imaged gel is completed by quantifying the bands using ImageQuant software and graphed with Prism. Each bar represents an average of three independent experiments (technical replicates) ± standard error as determined using Prism analysis software.

## 4. Results

### 4.1 Quantification of APE1 kinetic rates on replication fork mimic substrates

To study APE1 cleavage activity at stalled replication forks, we constructed three abasic DNA substrates that mimic various regions of a stalled replication fork. Specifically, the substrates each contain a single abasic site analog (tetrahydrofuran, THF) and were designed to represent varying degrees of interaction with a double stranded primer template junction (PTJ) (**Figure 2A**). Based on a structure of APE1 with duplex DNA (PDB 5DFI), the number of DNA nucleotides that interacts with APE1, termed its footprint, is about ten with APE1 centered at the abasic site^36, 39^. Utilizing this information, we designed the primer template junction (PTJ) substrate to contain an abasic site directly next to the double stranded 3′ end, requiring APE1 to interact with the double stranded junction (**Figure 2B**). The second substrate, a recessed PTJ substrate (Rec-PTJ), places the abasic site 7 nucleotides away from the double stranded 3′ end. In this case, the APE1 footprint is only contacting ssDNA, but is near the double stranded junction. Finally, we also designed a ssDNA substrate lacking any adjacent dsDNA, which represents APE1 contacting an abasic site within an extensive stretch of ssDNA upon replication fork stalling. Importantly, these substrates mimic common cellular situations by placing the abasic site right at a stalled replication fork, near a primer template junction, or within a stretch of ssDNA (**Figure 2**).

**Figure 2.**
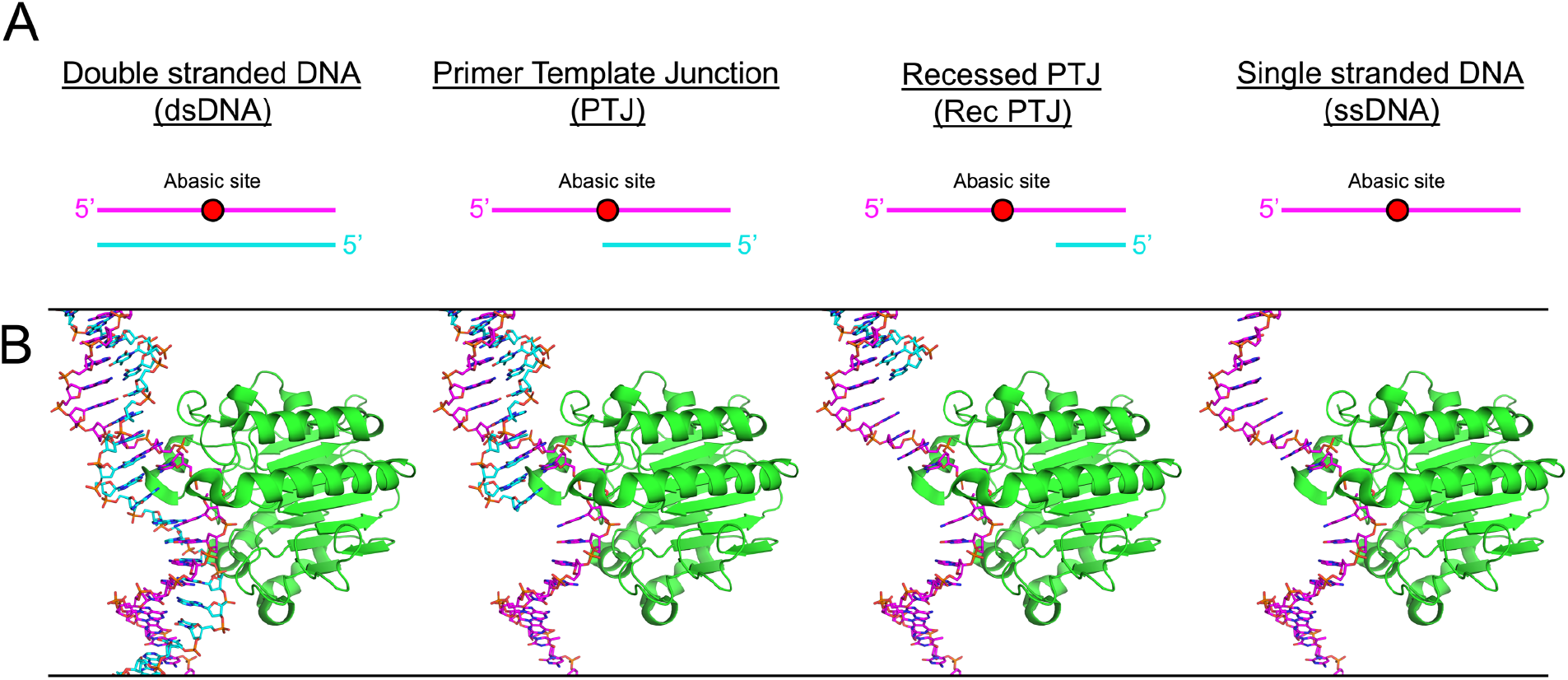
Replication fork mimic substrate design. (**A**) Cartoon representation of DNA substrates (represented as a pink and cyan lines) with an abasic site (represented as a red circle). (**B**) Structural models of PDB 5DFI with APE1 (shown as green ribbon) and DNA (shown as pink and cyan sticks).

To first determine whether APE1 can cleave these ssDNA substrates, we first utilized a qualitative product formation assay. To that end, we added full-length wild-type APE1 to each substrate and quenched the reaction with EDTA after one minute prior to separating the substrate and product bands on a denaturing Urea-PAGE gel. The formation of product was observed for each of the substrates, indicating APE1 indeed has cleavage activity on our replication fork mimic substrates (**Figure 3A**). Next, to quantitatively analyze the rate of APE1 cleavage activity, we performed pre-steady-state enzyme kinetics for the same set of ssDNA substrates. Under a multiple turnover kinetic regime (in which the concentration of DNA was in excess of APE1), the reaction produced a biphasic time course of product formation, consistent with what has been previously reported for APE1 endonuclease activity (**Figure 3B** and **3C**)^34^. This biphasic kinetic behavior is characterized by an initial burst phase corresponding to the first enzymatic turnover and the rate of DNA cleavage (k_obs_), and is followed by a rate-limiting steady-state phase, presumably corresponding to the rate of product release (k_ss_)^34^. For the PTJ substrate, the k_obs_ and k_ss_ were determined to be 10 ± 1 and 2.7 ± 0.21 sec^-1^, respectively (**Table 1**). Compared to rates determined for APE1 on the canonical dsDNA substrate, this represents a 12.9-fold decrease in catalysis and a 1.6-fold increase in product release for the PTJ substrate. For the recessed PTJ substrate, which has the abasic site further removed from the double stranded junction, k_obs_ was 30 ± 4 sec^-1^ and k_ss_ was 5.9 ± 0.32 sec^-1^, representing a 4.3-fold decrease in catalysis and 3.5-fold increase in product release (**Table 1**). For both PTJ substrates, the k_obs_ is <10-fold greater than the k_ss_ value, which makes accurate determination of k_obs_ difficult because the two phases are not easily differentiated during fitting.

**Figure 3.**
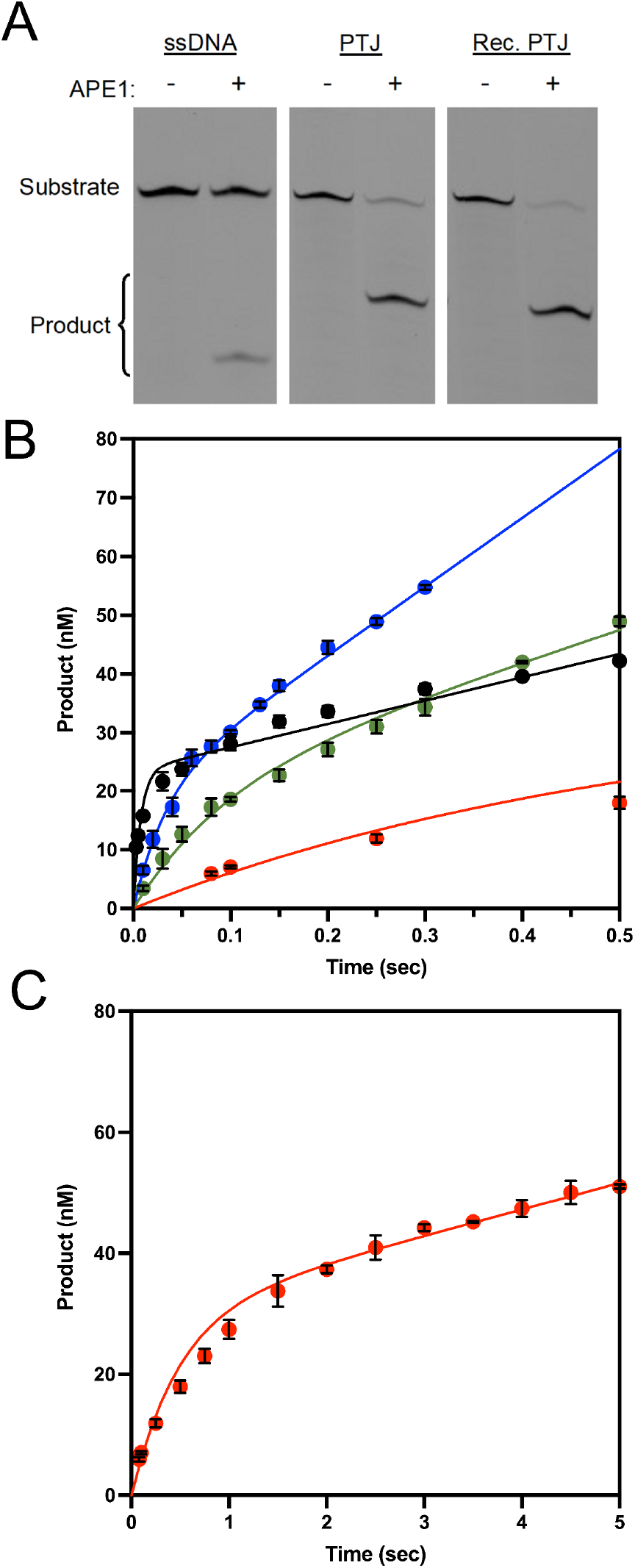
Cleavage activity of wild-type APE1 with PTJ and ssDNA substrates. (**A**) Product formation assay showing DNA bands separated on denaturing gel (also see Fig 3a source data 1 and 2) (**B**) Multiple turnover kinetic time courses of product formation for the reaction with line of best fit for each substrate; dsDNA (black), PTJ (green), recessed PTJ (blue), and ssDNA (red). (**C**) Extended time points for the ssDNA kinetic curve (red). All time points are the mean and standard deviation of three independent experiments with error bars (st. dev) shown. Where error bars are not seen, they are smaller than the data point.

**Table 1.** Pre-steady-state kinetic parameters of the wild-type APE1. Fold change is calculated from dsDNA values.

| Substrate | $k_{\text{obs}} \text{ (s}^{-1}\text{)}$ | $k_{\text{obs}}$<br>(fold change) | $k_{\text{ss}} \text{ (s}^{-1}\text{)}$ | $k_{\text{ss}}$<br>(fold change) |
| --- | --- | --- | --- | --- |
| dsDNA | $129 \pm 24$ | -- | $1.7 \pm 0.26$ | -- |
| PTJ | $10 \pm 1$ | $12.9 \pm 2.6 \downarrow$ | $2.7 \pm 0.21$ | $1.6 \pm 0.3 \uparrow$ |
| Rec PTJ | $30 \pm 4$ | $4.3 \pm 0.9 \downarrow$ | $5.9 \pm 0.32$ | $3.5 \pm 0.6 \uparrow$ |
| ssDNA | $2 \pm 0.2$ | $64.5 \pm 12.9 \downarrow$ | $0.15 \pm 0.018$ | $11.3 \pm 0.4 \downarrow$ |

To confirm the measured k_obs_ for both PTJ substrates, we performed additional single-turnover kinetic measurements. Consistent with the multiple-turnover measurements, the single turnover k_obs_ for the PTJ and Rec-PTJ substrates were determined to be 13.6 ± 0.4 and 25.7 ± 1.6 sec^-1^, respectively (**Figure S2**). Together, these results suggest that APE1 efficiently processes abasic sites at and near a primer template junction, with only a slight reduction in kinetic rate compared to dsDNA. Finally, for the ssDNA substrate, the k_obs_ was 2 ± 0.2 sec^-1^ and k_ss_ was 0.15 ± 0.018 sec^-1^, representing a 64.5- and 11.3-fold decrease in activity compared to dsDNA, respectively (**Table 1**). This is the most significant reduction in cleavage we observed and suggests that the presence of some form of duplex DNA, as with the PTJ, is preferred for optimal APE1 activity. Together, these kinetic data indicate that APE1 efficiently processes these replication fork mimic substrates, though the associated catalytic rates are reduced compared to the exceedingly fast APE1 endonuclease activity on duplex DNA.

To determine the binding affinity of APE1 for the replication fork mimic substrates, we performed electrophoretic mobility shift essays (EMSAs) in the presence of EDTA to inhibit catalysis. Quantification of the EMSAs yielded K_d,app_ values for the PTJ, Rec-PTJ, and ssDNA of 1.2 ± 0.1 nM, 0.7 ± 0.05 nM, and 22 ± 3 nM, respectively (**Figure 4A** and **4B**). Both PTJ and Rec-PTJ substrates had a similar effect on APE1 binding compared to dsDNA (K_d,app_ = 0.4 ± 0.1 nM), with a less than a 3-fold increase in their K_d,app_. In contrast, the affinity for ssDNA was decreased 55-fold compared to dsDNA. Nonetheless, an apparent binding affinity of 22 ± 3 nM still represents a very tight interaction of APE1 with ssDNA. Overall, these results demonstrate that APE1 binds to both the PTJ and the ssDNA substrates with a tight binding affinity.

**Figure 4.**
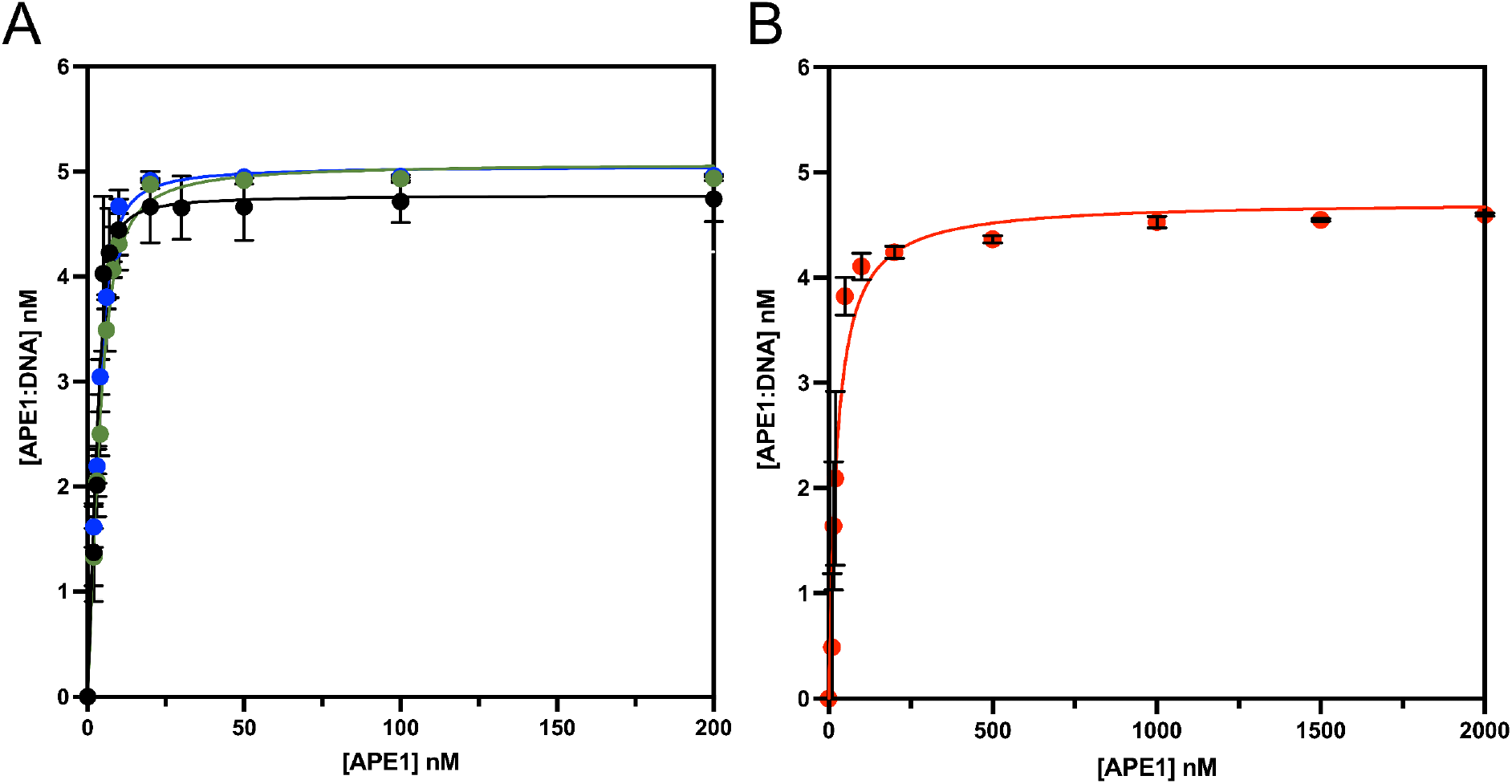
Binding of WT APE1 to PTJ and ssDNA substrates. Quantification of EMSA analysis. The line represents best fit to Equation 1 for (**A**) dsDNA (black), PTJ (green), and Recessed PTJ (blue). (**B**) Extended points for ssDNA curve (red). All points are shown as the mean of three independent experiments with error bars (st. dev.) shown. Where error bars are not seen, they are smaller than the data point.

### 4.2 Structural characterization of APE1 bound to a ssDNA substrate

To obtain structural insight into APE1 activity on ssDNA, we determined an X-ray crystal structure of APE1 bound to a ssDNA substrate (APE1:ssDNA). APE1:ssDNA crystals were generated using a 13-mer DNA oligonucleotide containing a centrally located abasic site analog. These crystals were generated in buffer conditions that included MgCl_2_, which supports APE1 catalysis, resulting in a product complex. Due to these conditions, we have been unable to capture the substrate complex. The resulting APE1:ssDNA crystals diffracted to a resolution of 2.0 Å in space group P65 (**Table S1**). The APE1:ssDNA complex structure represents a product complex, where APE1 catalysis has generated a nick in the DNA backbone. Importantly, as APE1 generated its product from substrate DNA, this signifies that APE1 is catalytically competent on the 13-mer ssDNA substrate. Moreover, as a result of APE1 cleavage, the downstream DNA sequence (5′ of the cleavage site) is not present in the structure of the APE1:ssDNA complex (**Figure 5A**). This is not surprising as APE1 has minimal interaction with DNA 5′ of the cleavage site.

**Figure 5.**
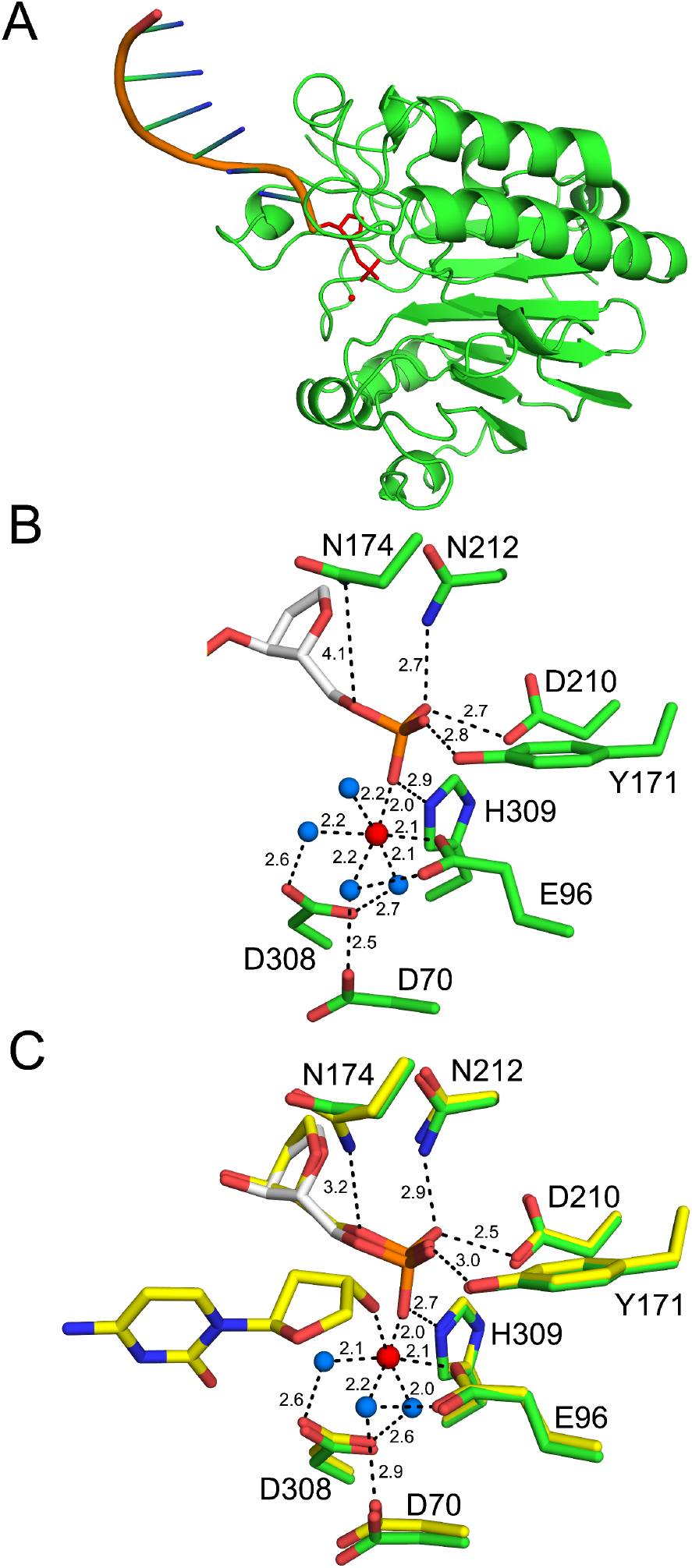
High-resolution structure of APE1: ssDNA product complex. (**A**) Overview of the APE1:ssDNA structure with APE1 (green, ribbons) and ssDNA (orange, cartoon). (**B**) A focused view of the APE1 active site with the THF (white) and key residues (green) shown in stick format (with distances in Å). The catalytic metal and active site waters are shown as red and blue spheres, respectively. (**C**) Overlay of ssDNA (green) with previous structure for WT APE1 with abasic dsDNA (yellow, PDB 5DFF). The catalytic metal, active site waters, and distances (Å) correspond to the dsDNA structure (PDB 5DFF).

The resulting APE1:ssDNA product complex reveals a well-formed APE1 active site, as seen previously for dsDNA structures (**Figure 5B**)^36, 39–40^. A single catalytic Mg^2+^ ion is directly coordinated by residue E96 and indirectly, through water-mediated interactions, by residues D70 and D308. Importantly, this Mg^2+^ is directly coordinated to the 5′-phosphate that is generated following cleavage. In the absence of the 3′-hydroxyl group (due to absence of the downstream DNA sequence in the crystal structure), an additional ordered water molecule coordinates the metal ion in the metal binding pocket. The non-bridging oxygens of the 5’-phosphate are coordinated by residues N212, D210, Y171, and H309, which stabilize the cleaved product complex.

Structural superimposition of the APE1:ssDNA product complex with the canonical APE1:dsDNA product complex (PDB 5DFF) highlights the structural similarities between the two complexes, where all the important catalytic residues are in nearly identical positions and maintain their respective interactions (**Figure 5C**). One minor difference is an ∼1 Å movement of residue N174 away from the abasic site 5′-phosphate. This movement disrupts the hydrogen bond between N174 and the bridging oxygen of the cleaved abasic site that is observed in the APE1:dsDNA structure. The subtle rearrangement of N174 likely reduces the stability of the abasic site and may contribute to the observed decrease in APE1 activity on ssDNA substrates. Together, this structure suggests APE1 utilizes the same active site residues and catalytic mechanism for processing ssDNA and dsDNA abasic sites.

### 4.3 Characterization of APE1 residue R177 during cleavage of abasic sites in ssDNA

In addition to the active site catalytic residues, we examined the roles of the APE1 DNA intercalating residues, R177. Structurally, these two residues intercalate between the two strands of double stranded DNA (**Figure 6A**). R177 acts as a surrogate base by intercalating into the major groove and forming base stacking interactions^39^. Additionally, R177 acts as a block between the abasic site and the orphan base, so they cannot hydrogen bond and the abasic site can be flipped out of the dsDNA into the active site for proper catalytic alignment. One unique property of APE1 processing ssDNA substrates is that there is no opposing non-damaged strand of DNA that must be held away from the abasic site strand during cleavage (**Figure 6B**).

**Figure 6.**
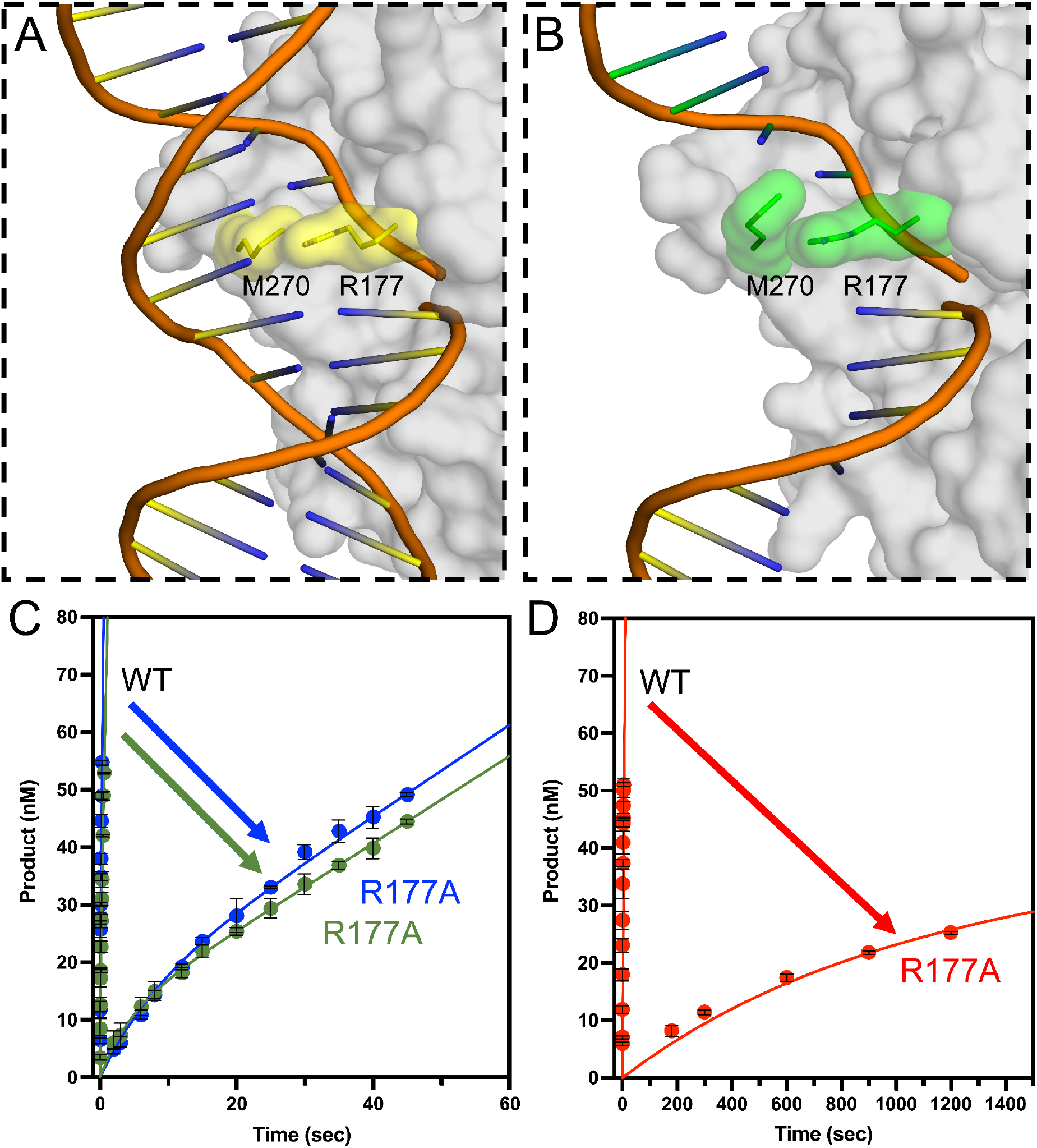
APE1 R177A mutant activity. (**A**) APE1 (grey, surface) bound to product dsDNA (orange, cartoon) highlighting DNA intercalating residues M270 and R177 (yellow, sticks) (PDB 5DFF). (**B**) APE1 (grey, surface) bound to ssDNA with downstream strand modeled in from PDB 5DFF highlighting M270 and R177 (green, sticks). Multiple turnover kinetic time courses of product formation for (**C**) PTJ (green) and Recessed PTJ (blue) and (**D**) ssDNA (red) substrates with line of best fit (in corresponding colors) compared to wild type. All time points are shown as the mean of three independent experiments with error bars (st. dev.) shown. Where error bars are not seen, they are smaller than the data point.

To determine if R177 is involved in processing of ssDNA substrates, we completed pre-steady-state kinetic analysis of the R177A mutant with each replication fork mimic substrate (**Figure 6C** and **6D**). For the PTJ substrate, the k_obs_ and k_ss_ were determined to be 0.23 ± 0.03 and 0.07 ± 0.003 sec^-1^, respectively. Compared to WT APE1, this represents a 43.5-fold decrease in catalysis and a 38.6-fold decrease in product release. For the recessed PTJ substrate, which has the abasic site further removed from the double stranded junction, k_obs_ was 0.12 ± 0.015 sec^-1^ and k_ss_ was 0.06 ± 0.006 sec^-1^, representing a 250-fold decrease in catalysis and 98.3-fold decrease in product release (**Figure 6C** and **Table 2**). Finally, for the ssDNA substrate, k_obs_ was determined to 0.001 ± 0.0001 and the k_ss_ 0.0001 ± 0.000008 sec^-1^, representing a significant decrease in activity with 2,000- and 1,500-fold decreases, respectively (**Figure 6D** and **Table 2**). Together, these data indicate that R177 serves an important function in the APE1 catalysis of ssDNA that is unique from its activity on dsDNA. We propose that R177A stabilizes the bases flanking the abasic site to provide stability to the base stacking of the DNA in the absence of a complimentary DNA strand.

**Table 2.** Pre-steady-state kinetic parameters of the R177A APE1. Wild-type values are the same as those shown in Table 1 for comparison to R177A values. Fold change is calculated for each substrate using the corresponding wild-type value.

| APE1 | Substrate | $k_{obs} \text{ (s}^{-1}\text{)}$ | $k_{obs}$<br>(fold change) | $k_{ss} \text{ (s}^{-1}\text{)}$ | $k_{ss}$<br>(fold change) |
| --- | --- | --- | --- | --- | --- |
| WT | PTJ | $10 \pm 1$ | -- | $2.7 \pm 0.21$ | -- |
| R177A | PTJ | $0.23 \pm 0.03$ | $43.5 \pm 6.9 \downarrow$ | $0.07 \pm 0.003$ | $38.6 \pm 3.4 \downarrow$ |
| WT | Rec PTJ | $30 \pm 4$ | -- | $5.9 \pm 0.32$ | -- |
| R177A | Rec PTJ | $0.12 \pm 0.015$ | $250 \pm 45.5 \downarrow$ | $0.06 \pm 0.006$ | $98.3 \pm 2.5 \downarrow$ |
| WT | ssDNA | $2 \pm 0.2$ | -- | $0.15 \pm 0.018$ | -- |
| R177A | ssDNA | $0.001 \pm 0.0001$ | $2000 \pm 280 \downarrow$ | $0.0001 \pm 0.000008$ | $1500 \pm 216 \downarrow$ |

As seen in the WT kinetics for both PTJ substates, k_obs_ is <10-fold greater than the k_ss_ value, which makes accurate determination of k_obs_ difficult. To confirm the reported k_obs_ for both PTJ substrates, we preformed additional single-turnover kinetic measurements with the R177A mutant. As observed previously with abasic dsDNA single-turnover kinetics, curves are best fit with a double exponential equation, indicating the presence of two distinct cleavage rates^34^. Here, we will focus our discussion on the rate determined for the major population for each substrate. The single turnover k_obs_ for the PTJ and Rec-PTJ substrates were determined to be 0.05 ± 0.004 and 0.06 ± 0.006 sec^-1^, respectively (**Figure S3**). In this case, since the single turnover k_obs_ values are very similar to the determined k_ss_ values, single turnover measurements provide a more accurate rate than multiple turnover measurements. However, both values emphasize the drastic reduction in activity observed for the R177 mutant from WT rates, highlighting this residue’s importance.

### 4.4 APE1 exchange for RPA on abasic PTJ substrates

At a stalled replication fork, exposed single stranded DNA is quickly bound by the major ssDNA binding protein, RPA. To study the interplay between RPA and APE1 on our PTJ substrates, we employed a novel FRET assay. This assay utilizes Cy5-labeled RPA and Cy3-labeled PTJ DNA substrates that contain an abasic site either directly next to the junction (PTJ) or 7 nucleotides away (Rec-PTJ), as used in the kinetic assays presented previously. The primer has a Cy3 FRET donor attached at the 3′ terminus. The length (33 nt) of the 5′ ssDNA overhang accommodates at least one RPA heterotrimeric complex^41^(**Figure 7A**). The RPA complex is labeled with a Cy5 FRET acceptor at position 107 of the RPA32 subunit, which lies within OB-fold D^33^ (**Figure 7B**). RPA resides on the 5′ ssDNA overhang with specific polarity such that the Cy5-labeled DBD-D is oriented towards the Cy3-labeled PTJ^33^ (**Figure 7C**). With RPA titration, the FRET signal increases linearly to a plateau and then remains constant, indicating that both DNA substrates can be saturated with Cy5-RPA (**Figure S4** and **S5**). In the APE1 experimental samples (**Figure 8A**), Cy3-labeled PTJ is pre-saturated with Cy5-labeled RPA, yielding a high FRET state, and the resultant mixture is then titrated with catalytically dead APE1 (i.e., APE1_Dead_). E_FRET_ is calculated after each addition of APE1_Dead_. Under the conditions of the assay, a decrease in E_FRET_ indicates that APE1_Dead_ engages the DNA and either; 1) alters the configuration of the RPA complex on DNA such that the distance between the FRET dyes increases (yielding an intermediate FRET state); 2) displaces RPA from the DNA (yielding a low FRET state) or; 3) a combination of both possibilities. For both the PTJ and Rec-PTJ substrates, formation of the DNA-RPA complex is indicated by a robust FRET signal before APE1_Dead_ is added.

**Figure 7.**
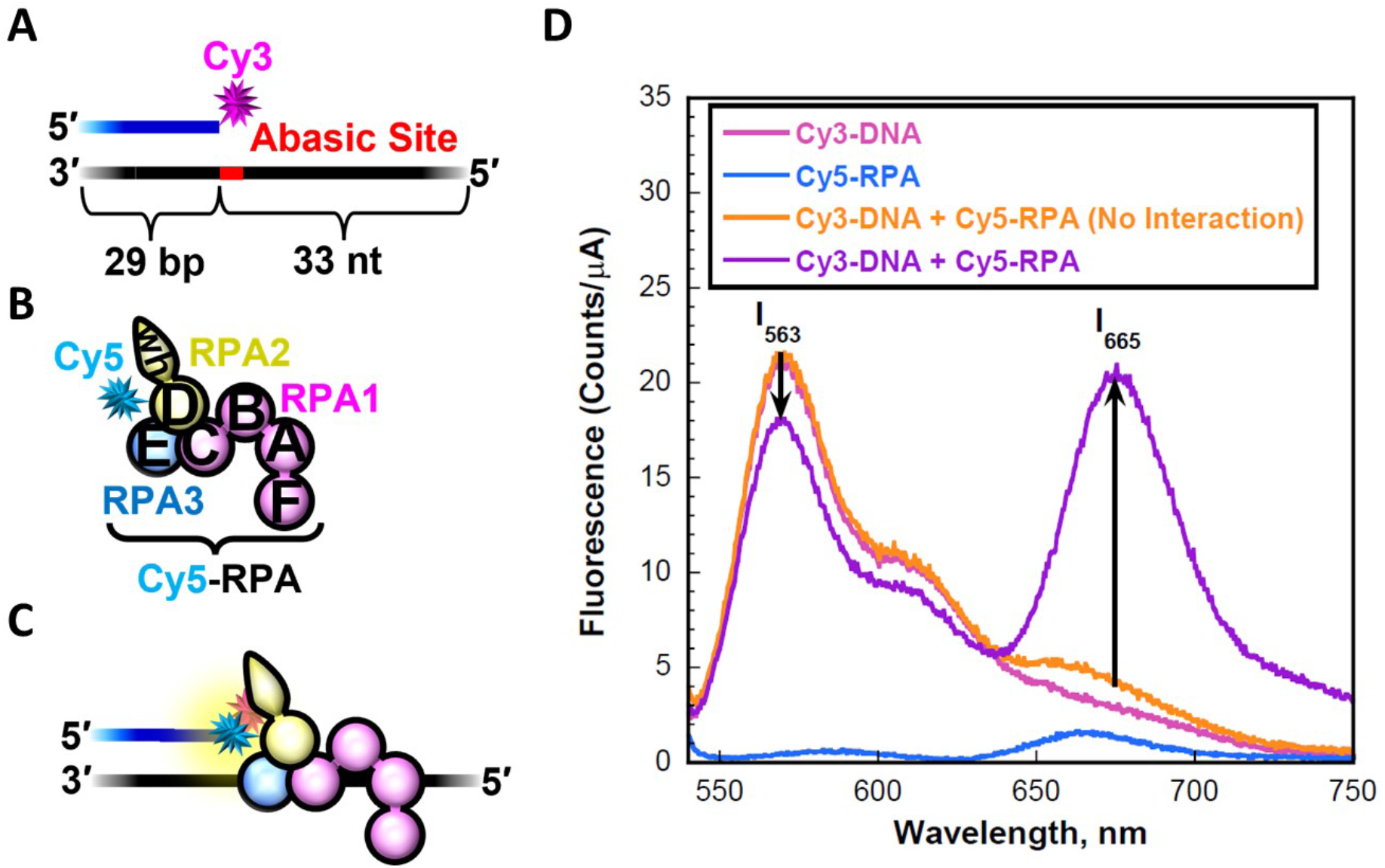
RPA engaging a PTJ that contains an abasic site. (**A**) Schematic representation of the 3′ Cy3-labeled PTJ substrate (**Figure S1**) (**B**) Schematic representation of Cy5-labeled RPA. The RPA subunits are color-coded (RPA1 in pink, RPA2 in yellow, and RPA3 in blue) and depicted to illustrate the OB-folds (A –E) and winged-helix (wh) domain. The RPA complex is labeled with a Cy5 FRET acceptor at residue 101 of the RPA32 subunit. This position lies within OB-fold D. (**C**) Schematic representation of RPA interactions at a PTJ. RPA interacts with the 5′ ssDNA overhang in an orientation-specific manner such that the Cy5-labeled RPA32 subunit is oriented towards the Cy3-labeled PTJ. (**D**) Fluorescence emission spectra (540 – 750 nm) obtained by exciting the Cy3-labeled PTJ DNA with 514 nm light. The fluorescence emission intensities (*I*) at 665 nm (Cy5 FRET acceptor fluorescence emission maximum, *I_665_*) and 563 nm (Cy3 FRET donor fluorescence emission maximum, *I_563_*) are indicated. Cy5-labeled RPA can be excited via FRET from Cy3 only when the two fluorophores are in close proximity (i.e., < 10 nm). This is indicated by an increase in I_665_ and a concomitant decrease in *I_563_*. The predicted spectrum for no interaction between RPA and DNA is determined by adding the spectrums of the individual components.

**Figure 8.**
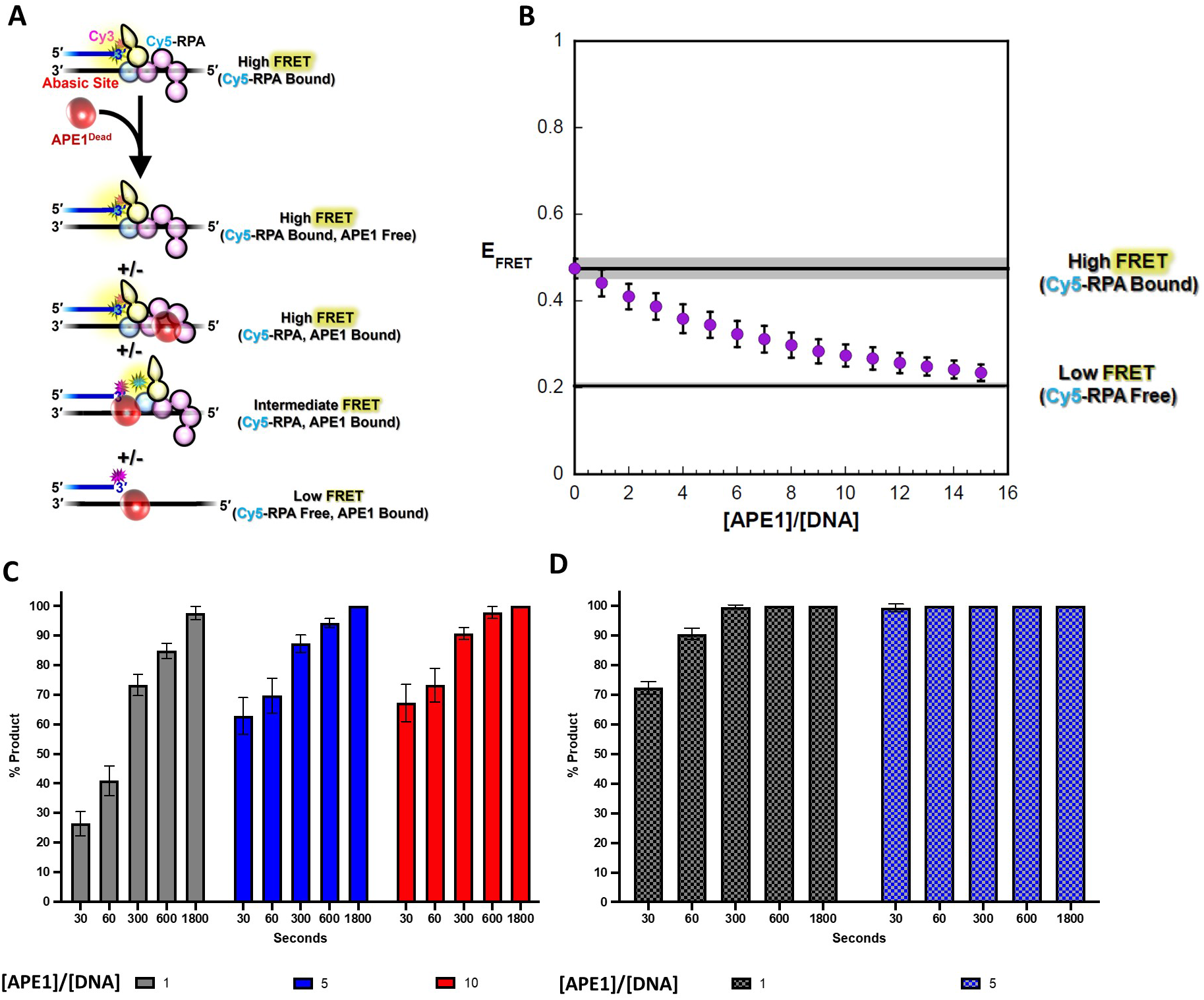
RPA/APE1 exchange on the Rec-PTJ substrate. (**A**) Schematic representation of the FRET experiment. A 3′ Cy3-labeled Rec-PTJ DNA containing an abasic site 7 nt downstream of the PTJ (25 nM, **Figure S1**) is pre-saturated with Cy5-labeled RPA, yielding a high FRET state, (**Figure S3**) and the resultant mixture is then titrated with catalytically dead APE1 (i.e., APE1_Dead_). E_FRET_ is calculated after each addition of protein. The ssDNA downstream of the PTJ (33 nt) can accommodate a single Cy5-RPA or 3 APE1_Dead_. Possible protein•DNA complexes and their expected E_FRET_ values are depicted. (**B**) E_FRET_ values are plotted as a function of [APE1_Dead_]/[DNA] ratio and each data point represents the mean ± S.E.M. of at least three independent measurements. The E_FRET_ value observed in the absence of APE1_Dead_ (0.475 ± 0.0227) is extrapolated to the axis limits as a flat black line (with S.E.M. displayed in grey) to indicate the high FRET state where the DNA substrate is devoid of APE1_Dead_. The predicted E_FRET_ value for no interaction between RPA and DNA (0.204 ± 0.00394) is displayed as a flat black line (with S.E.M. displayed in grey) to indicate the low FRET state where the DNA substrate is devoid of RPA. The observed FRET decreases to within experimental error of the low FRET state over the range of APE1_Dead_ additions, indicating that APE1_Dead_ completely exchanges for RPA. (**C**) Ape1 product formation (%) values over time in the presence of pre-saturated RPA on Rec-PTJ substrate. **(D)** Ape1 product formation (%) in absence of RPA on Rec-PTJ substrate. [APE1_Dead_]/[DNA] ratio of 1, 5, and 10 indicated by color as grey, red and blue, respectively with each bar representing the mean ± S.E.M. of at least three independent measurements.

For the Rec-PTJ substrate (**Figure 8B**), which contains an abasic site 7 nucleotides downstream of the PTJ, a robust FRET signal is observed prior to the addition of APE1_Dead_ (High FRET, Cy5-RPA Bound). Over the course of the APE1_Dead_ titration, the observed E_FRET_ values decrease to within experimental error of the low FRET state where the DNA substrate is devoid of RPA. This indicates complete exchange of RPA for APE1_Dead_ on the Rec-PTJ DNA substrate. The intermediate FRET values observed during the titration may represent RPA binding in different conformations before its complete exchange at the high APE1 concentrations. In the FRET assay, it was necessary to use catalytically dead APE1 to prevent cleavage from occurring, however it is also of interest to determine if APE1 is able to cut the DNA substrate under these experimental conditions. To test APE1 activity, we analyzed APE1 product formation under the same experimental conditions and protein ratios as the FRET experiments. The product formation assay demonstrates that at a [APE1]/[DNA] = 1, about 25% product was generated after only 30 seconds and increasing product was generated over time (**Figure 8C**). As expected, we observe an APE1 concentration-dependent increase in activity. Compared to the no RPA control, this represents a decrease in APE1 activity in presence of RPA, which is consistent with previous publications reporting RPA suppressing APE1 activity^26^ (**Figure 8D**). However, this data demonstrates that APE1 maintains robust cleavage activity in the presence of RPA.

For the PTJ substrate, where the PTJ directly abuts an abasic site, the observed E_FRET_ values do not change over the course of the APE1_Dead_ titration. This indicates that APE1_Dead_ does not exchange for RPA nor promote a long-lived conformational re-arrangement of RPA that increases the distance between the FRET dyes (**Figure 9A** and **9B**). However, it should be noted that we cannot rule out that APE1_Dead_ engages RPA-bound DNA in a manner that does not alter the distance between the FRET dyes. Thus, at least DBD-D (and likely DBD-E) of the RPA complex remain stably bound to the PTJ even at very high concentrations of APE1_Dead_. Remarkably, product formation assays reveal that APE1 is able to cleave the abasic DNA under these conditions. As seen for the Rec-PTJ, we observe increasing APE1 activity over time and with increasing APE1 concentrations (**Figure 9C** and **9D**). This data demonstrates that APE1 is active on the abasic PTJ substrate with RPA still bound to the DNA. It is likely that APE1 is cutting both the Rec-PTJ and the PTJ substrates while RPA is bound, however in the Rec-PTJ the position of the abasic site is allowing APE1 to kick RPA off the substrate at very high APE1 concentrations.

**Figure 9.**
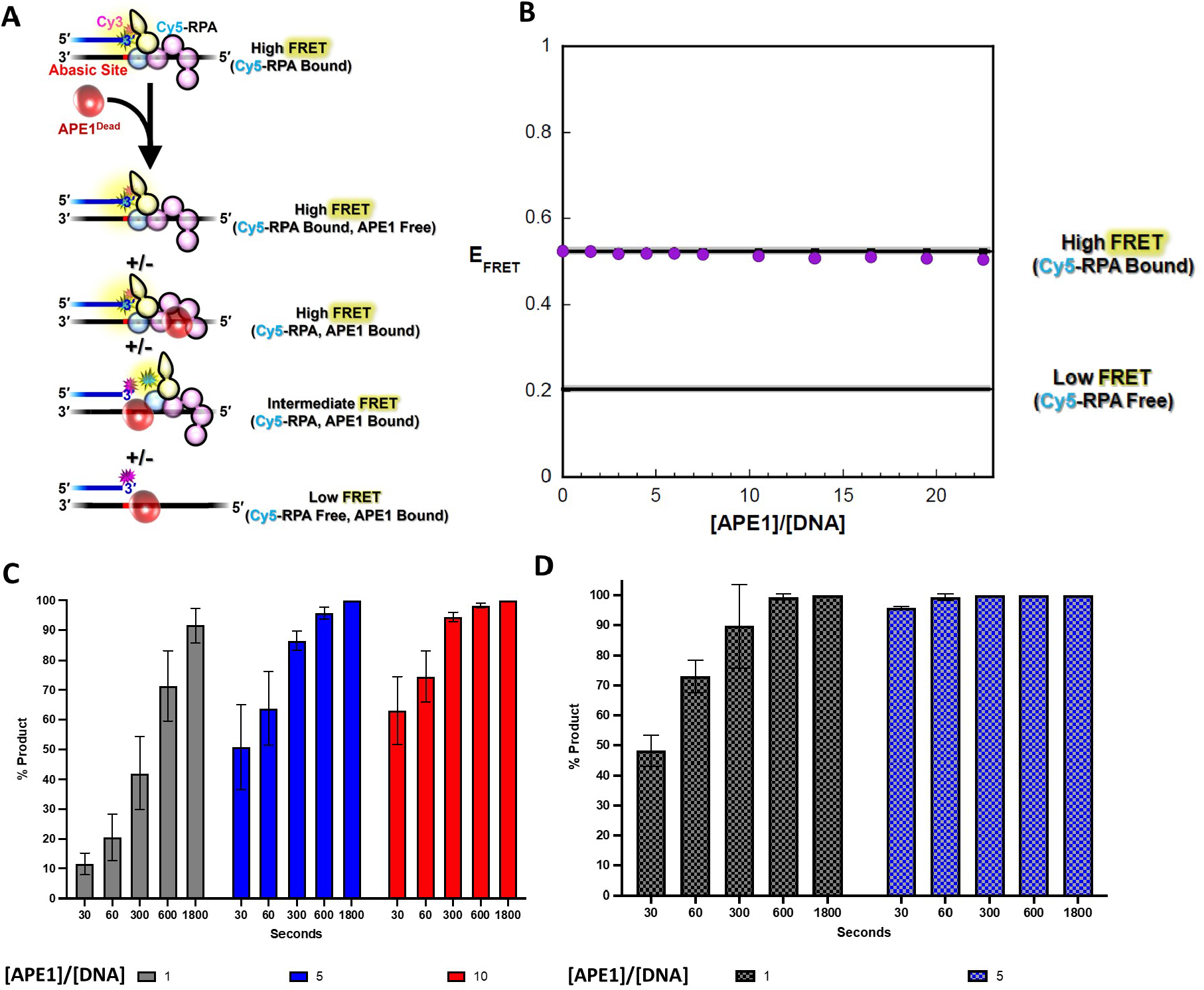
RPA/APE1 exchange on the PTJ substrate. (**A**) Schematic representation of the FRET experiment. A 3′ Cy3-labeled PTJ DNA containing an abasic site at the PTJ (25 nM, Figure S1) is pre-saturated with Cy5-labeled RPA (**Figure S2**), yielding a high FRET state, and the resultant mixture is then titrated with catalytically dead APE1 (i.e., APE1_Dead_). E_FRET_ is calculated after each addition of protein. The ssDNA downstream of the PTJ (33 nt) can accommodate a single Cy5-RPA or 3 APE1_Dead_. Possible protein•DNA complexes and their expected E_FRET_ values are depicted. (B) E_FRET_ values are plotted as a function of [APE1Dead]/[DNA] ratio and each data point represents the mean ± S.E.M. of four independent measurements. The E_FRET_ value observed in the absence of APE1_Dead_ (0.524 ± 0.00608) is extrapolated to the axis limits as a flat black line (with S.E.M. displayed in grey) to indicate the high FRET state where the DNA substrate is devoid of APE1_Dead_. The predicted E_FRET_ value for no interaction between RPA and DNA (0.205 ± 0.00538) is displayed as a flat black line (with S.E.M. displayed in grey) to indicate the low FRET state where the DNA substrate is devoid of RPA. The observed FRET does not decrease over the range of protein additions, indicating that RPA is not exchanged for APE1. (**C**) Ape1 product formation (%) values over time in the presence of pre-saturated RPA on PTJ substrate. **(D)** Ape1 product formation (%) in absence of RPA on PTJ substrate. [APE1_Dead_]/[DNA] ratio of 1, 5, and 10 indicated by color as grey, red and blue, respectively with each bar representing the mean ± S.E.M. of at least three independent measurements.

To account for the position of the abasic site in the PTJ substrates, we used competitor DNA to determine the relative binding affinity of RPA for each substrate. When the unlabeled competitor DNA is identical to the Cy3-labeled DNA substrate on which the FRET complex is formed, it is expected that 50% inhibition is observed when the concentration of competitor DNA is equal to the concentration of the substrate DNA, i.e., [Competitor DNA]/[DNA]_Total_ = 0.50. Using Cy3-labeled Rec-PTJ and unlabeled competitor Rec-PTJ DNA, the observed [Competitor DNA]/[DNA]_Total_ is equal to 0.487 ± 5.61 x 10^-3^, confirming the validity of the approach and confirming that the Cy3 label at the PTJ has no effect on the binding of RPA to the DNA substrates (**Figure S6**). When the competitor DNA is instead the PTJ DNA substrate (contains an abasic site at the PTJ) the observed [Competitor DNA]/[DNA]_Total_ decreases slightly to 0.413 ± 0.0447, indicating that RPA has a slightly higher relative affinity for a PTJ substrate compared a Rec-PTJ substrate (**Figure S6**). However, this slight difference in affinities does not account for the drastic difference in RPA/APE1 exchange for the PTJ and Rec-PTJ DNA substrates (**Figure 8B** and **9B**). Hence, the observed differences in exchange of APE1 for RPA on the PTJ DNA substrates are likely attributed to differential dynamics of RPA on the abasic site-containing PTJ DNAs.

## 5. Discussion

### 5.1 APE1 cleavage of ssDNA substrates

Previous work has demonstrated APE1 cleavage activity on several complex structures, such as primer-template duplexes, bubbles, and forks^25–26, 42–43^. Here, we advance our understanding of this APE1 activity using pre-steady-state kinetics, binding, and X-ray crystallography to determine how APE1 processes ssDNA abasic sites in replication fork mimic substrates. Data demonstrate that APE1 binds these substrates with high affinity and cleaves with only moderately reduced activity compared to canonical APE1 cleavage of double stranded substrates. The kinetic values of APE1 cleavage are well within the range of reported rates for other APE1 substates, as well as other DNA repair enzyme activities^31, 44–49^. It has been previously observed that some form of DNA secondary structure is preferred for APE1 activity, but that proximity of the abasic site influences APE1 incision efficiency^26^. Our data supports this observation, as APE1 cleavage efficiency is affected by the position of the abasic site compared to the double stranded PTJ.

### 5.2 R177 is critical to APE1 ssDNA cleavage activity

Since the first structural observations of APE1 residue R177 intercalating dsDNA, this residue have been of interest in the APE1 field^36^. Today, R177 has been implicated in several functions in the APE1 catalytic mechanism including product release, protein-DNA interactions, and substrate specificity^31, 36, 39, 50–52^. Mutation of R177 to alanine results in only subtle increases in kinetic rate for both the AP-endonuclease and proofreading exonuclease activities^31, 39^. However, a recent publication reported that R177A decreased APE1 activity when processing a 3’-8oxoG lesion, indicating R177 plays an important role in mechanism of 8oxoG processing by APE1^53^.

Previous descriptions all include APE1 interacting with both strands of dsDNA substates, prompting our interest in the function of these residues in the context of ssDNA substrates. The R177A mutation resulted in dramatically reduced APE1 kinetic rates on all three of the replication fork mimic substrates. In fact, kinetic data indicates that the R177 side chain becomes more critical for APE1 activity as the flanking dsDNA decreases (as demonstrated by lowest rate for ssDNA, **Table 2**). Based on the data presented here, we propose that R177A plays a role in stabilizing the bases flanking the abasic site to provide stability to the base stacking of the DNA in the absence of a complimentary DNA strand. Additionally, it is likely that R177 is playing a “space filling role” acting as a physical block to hold the abasic site out of the double helix and into the active site for cleavage, similar to the role described as the “RM bridge”^52^. These experiments, along with structural observations, provide insight into why R177 may have a more critical role in APE1 cleavage of ssDNA over its modest role observed for canonical APE1 function processing dsDNA.

### 5.3 APE1:DNA interactions

Although there is currently no structure of APE1 bound to a primer template junction substrate, structural data presented here and published previously provide insight into the expected interactions between APE1 and a PTJ. APE1 interactions with double stranded DNA have been classified into two regions or “hot spots” (**Figure 10A**)^52^. These two regions are (1) with the abasic strand backbone flanking the cleavage site and (2) with the opposing nondamaged strand backbone downstream (3’) of the orphan base (**Figure 10A**). The latter of these interaction hot spots would be lost in an APE1-PTJ interaction, as the 3’ region of the nondamaged strand would be missing from a PTJ structure (**Figure 10A**, cyan bracket). Alternatively, when interacting with a ssDNA-dsDNA PTJ, APE1 may utilize a novel, and currently unknown, mode of binding to facilitate catalysis.

**Figure 10.**
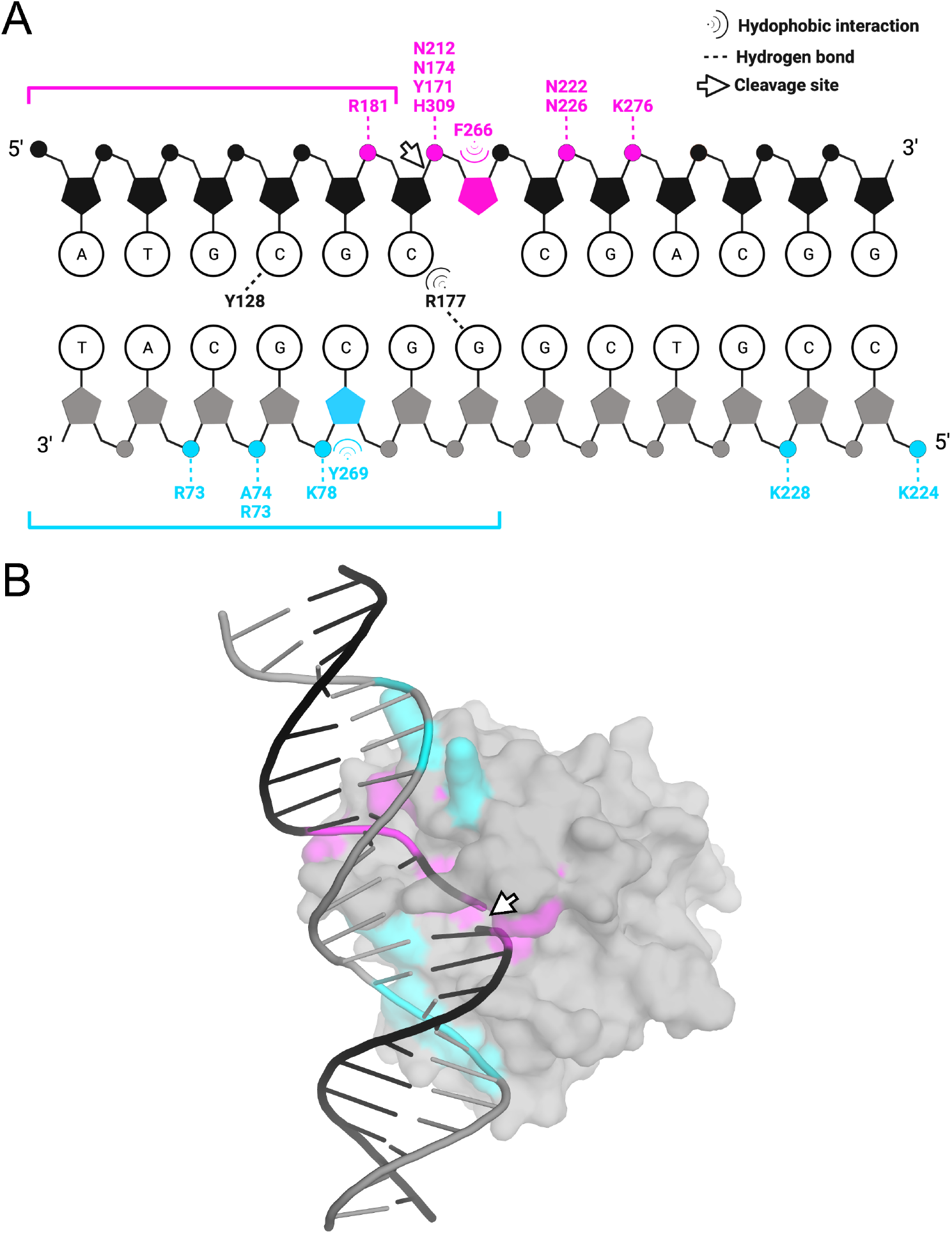
APE1 interactions with abasic dsDNA. (**A**) Schematic of the interactions between wild-type APE1 and abasic dsDNA. Interactions with abasic and opposing nondamaged strands are shown in pink and cyan, respectively. The pink bracket highlights the 5’ downstream region of the abasic strand that is absent from the APE1:ssDNA product complex crystal structure. The cyan bracket highlights the region of the nondamaged strand that would be absent in a PTJ structure. Interactions were determined using the protein–ligand interaction profiler, PLIP, available at https://plip-tool.biotec.tu-dresden.de with hydrogen bonds limited at 3.8 Å^54^. Figure created with BioRender.com. (**B**) Overview of APE1:AP-dsDNA structure with APE1 (grey, surface) bound to dsDNA (black and grey, cartoon). The DNA and interacting residues (as shown in panel **A**) are highlighted in pink or cyan, based on interaction with the abasic or opposing nondamaged strand, respectively. Cleavage site is denoted with a white arrow (PDB 5DFF).

The interaction schematic highlights that the majority of APE1 interactions are with the abasic strand of DNA (**Figure 10B**). In this way, APE1 approaches the DNA from the damaged side and “sits” on the abasic strand. Unlike polymerases, which use the templating strand for insertion, APE1 does not rely on the opposing strand for activity, and is thus capable of cleaving ssDNA substrates, as demonstrated by the data presented in this study. Furthermore, APE1 does not make significant contacts with the damaged strand 5’ of the active site (**Figure 10A**, pink bracket), as demonstrated in substrate complex (PDB 5DFI), product complex (PDB 5DFF), as well as the loss of this strand in the APE1:ssDNA product complex structure presented here (**Figure 5A**)^54^. In a cellular context, this would present a biological problem if APE1 cuts the backbone of a ssDNA substrate and immediately loses contact with this portion of the DNA. In this way, APE1 cleavage could instantly collapse the replication fork, potentially even while APE1 is still bound. One hypothesis is that fragile repair intermediates are protected during repair through substrate channeling among repair enzymes. Therefore, other enzymes are likely needed to protect such a repair intermediate for other pathways that are necessary for repair. Alternatively, APE1 cleavage of ssDNA may be very deleterious to the cell, and therefore depend heavily on proteins to protect abasic ssDNA from APE1 cleavage.

### 5.4 Interplay between APE1 and RPA

During replication fork stalling, the DNA helicase continues to unwind the dsDNA generating large stretches of excess ssDNA which are then coated by RPA^15–16, 55–57^. By binding transiently exposed ssDNA, RPA serves as a hub protein to coordinate DNA replication, as well as other cellular processes^58–59^. RPA binds to ssDNA with a sub-nanomolar affinity; however, the RPA-ssDNA complex is dynamic, and several studies have suggested the RPA DNA-binding domains (DBDs) undergo microscopic dissociations from the ssDNA^41, 60–61^. These microscopic dissociations from the DNA are thought to enable lower affinity DNA-binding proteins to displace and/or remodel RPA.

Here, we observe that APE1 is able to cleave the phosphodiester backbone of an abasic PTJ substrate while RPA remains bound. In this way, APE1 and RPA are accommodated on the same DNA substrate. Furthermore, when the abasic site is positioned further from the PTJ in the ssDNA region, APE1 is able to displace RPA at high concentrations. This is consistent with a dynamic RPA-DNA complex allowing APE1 to interact with exposed DNA or during micro dissociation events. While RPA does have a large interaction “footprint” with the DNA (18-22 nt), structural remodeling or basal diffusion of RPA exposes small stretches of ssDNA and likely allows APE1 to gain access to a lesion for rapid cleavage. Data reported here supports a dynamic RPA-DNA complex which allows access to other DNA repair factors, such as APE1. This model is further supported by recent work that demonstrated RPA DBD-A and DBD-B are highly dynamic and that the RPA-ssDNA complex exists in at least four distinct conformations that each allow varying access to the ssDNA^62^. This is consistent with data showing that the interaction of RPA with the uracil DNA glycosylase (UNG) stimulates the excision of uracil from RPA-coated ssDNA via increased accessibility of ssDNA^9^.

Importantly, APE1 cleavage of ssDNA containing abasic sites at the replication fork would have significant implications for genomic stability, as it would generate a double stranded break that could collapse the replication fork. This suggests that cellular mechanisms likely exist to prevent APE1 cleavage of ssDNA or to protect the nicked ssDNA following APE1 cleavage. While RPA does not directly interact with APE1, it has been reported to act as an inhibitor of APE1 ssDNA cleavage activity through an unknown mechanism^26^. Consistently, we observed a reduction in APE1 product formation in the presence RPA. However, it is important to consider the abundance of cellular APE1 when choosing experimental conditions. We observed that even with RPA pre-saturated on the DNA, APE1 has a robust cleavage activity over a range of APE1 concentrations. It is possible that RPA-coated ssDNA could inhibit APE1 binding or alter ssDNA structural elements necessary for APE1 activity. However, the kinetic rates determined here identify that APE1 rapidly cleaves abasic sites in replication fork mimic substrates. Further work is needed to elucidate the complex interplay between APE1 and RPA at replication forks containing abasic sites.

## Acknowledgements

This research was funded by National Institute of Environmental Health Sciences [R01ES029203 to BDF] and the National Institute of General Medical Science [R01GM133967 and R01GM130746 to EA,1R35GM147238-01 to MH] of the National Institutes of Health, and the American Heart Association [20PRE35210752 to NMH]. AUC experiments to characterize the fluorescent human RPA were supported by S10OD030343 to EA.

## Author Contributions

B.D.F. and N.M.H. designed the study. N.M.H., J.L.N., and V.K. expressed and purified the proteins. T.H.K. collected the crystal structure. N.M.H performed kinetic and binding data acquisition, X-ray crystallographic structure refinement/deposition. M.H. designed the FRET experiments. J.L.N. and V.K. performed the FRET experiments. B.D.F., M.H., E.A., N.M.H analyzed data and wrote the manuscript.

## Data Availability

Atomic coordinates and structure factors for the reported crystal structure have been deposited with the Protein Data bank under accession number 7TR7.

## Supplementary Figures

**Figure S1.**
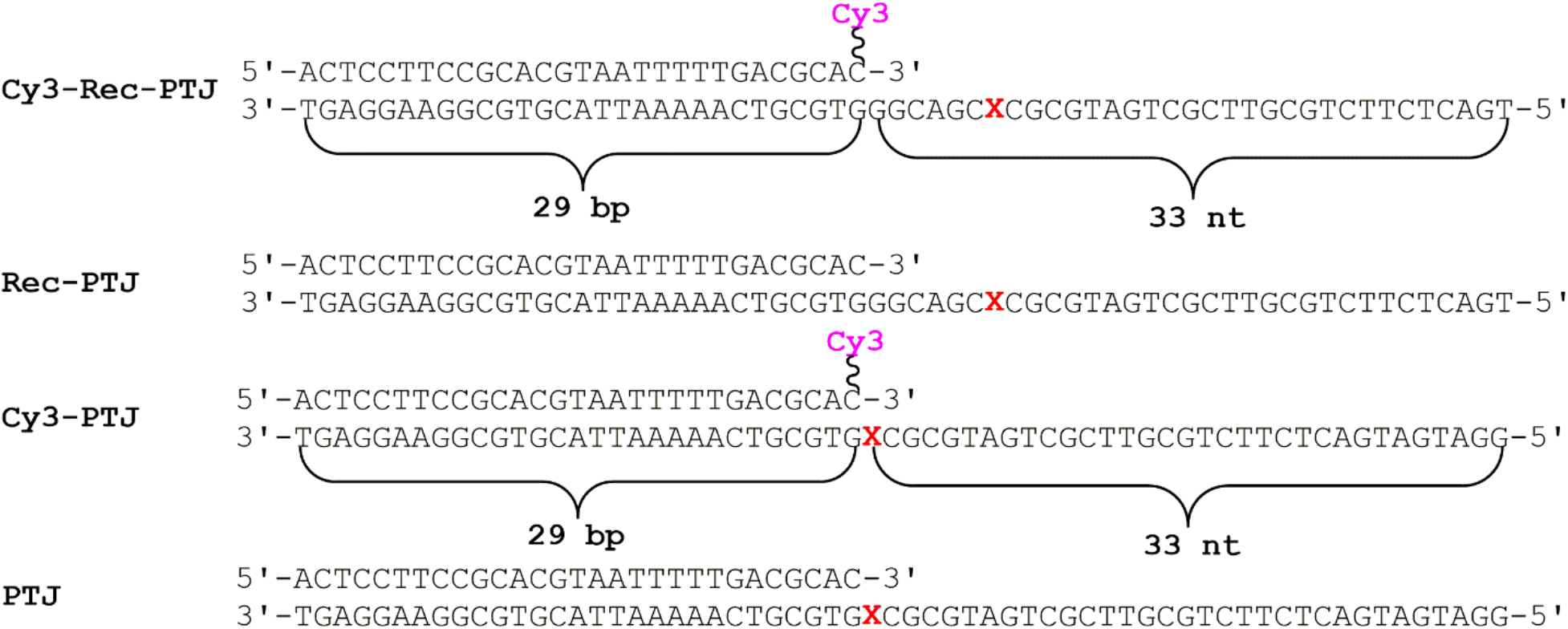
PTJ DNA substrates utilized in this study. The sequences and lengths (29 bp) of the double strand DNA (dsDNA) regions are identical. X denotes an abasic site. The ssDNA regions adjacent to the 3’-end of the Primer-Template junctions (PTJ) accommodate at least 1 RPA molecule^41^.

**Figure S2.**
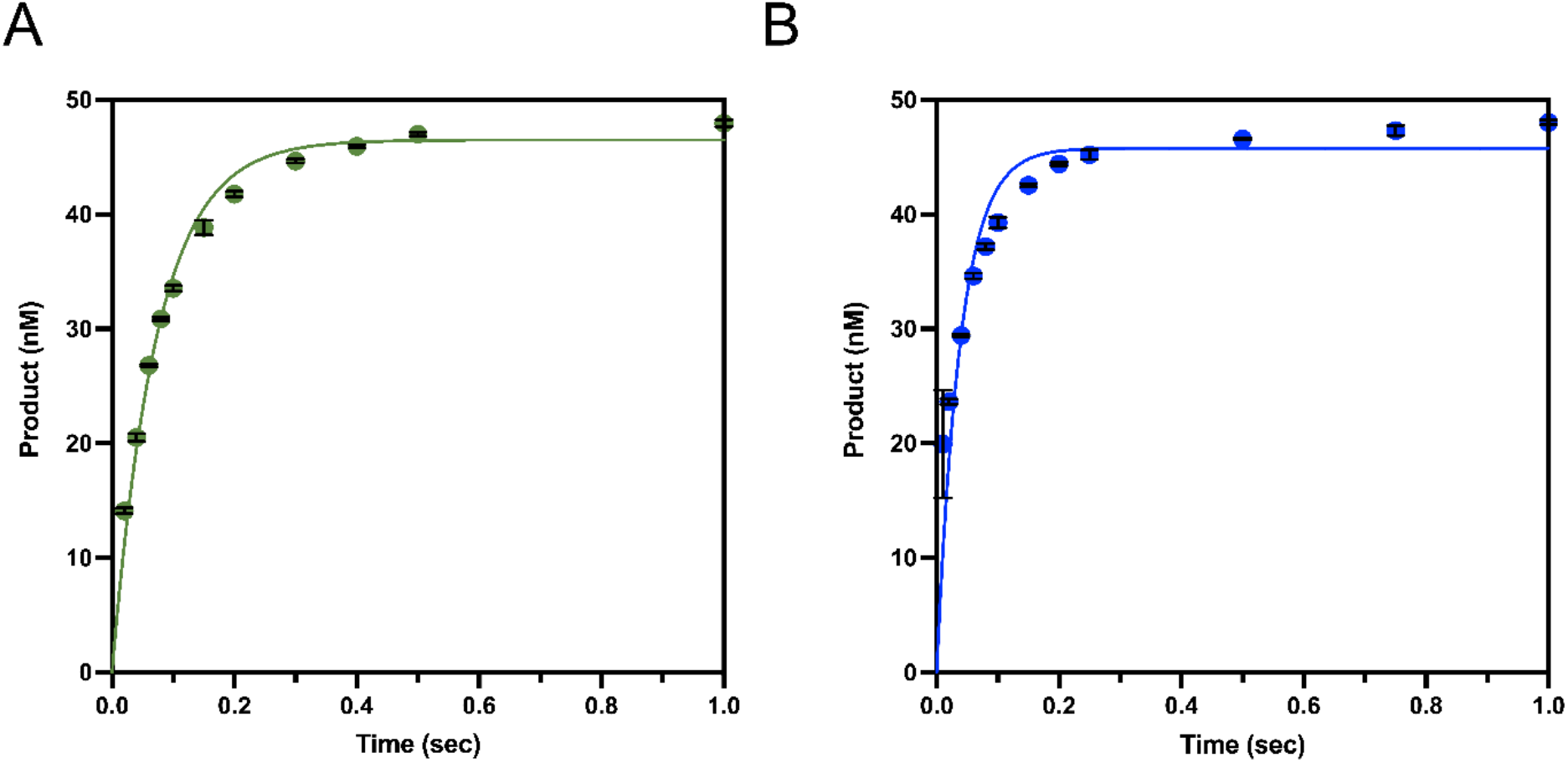
Single turnover kinetic time courses. Single turnover kinetic time courses for WT APE1 with (**A**) PTJ (green) and (**B**) Rec-PTJ (blue). All time points are shown as the mean of three independent experiments with error bars (st. dev.) shown. Where error bars are not seen, they are smaller than the data point.

**Table S1.**
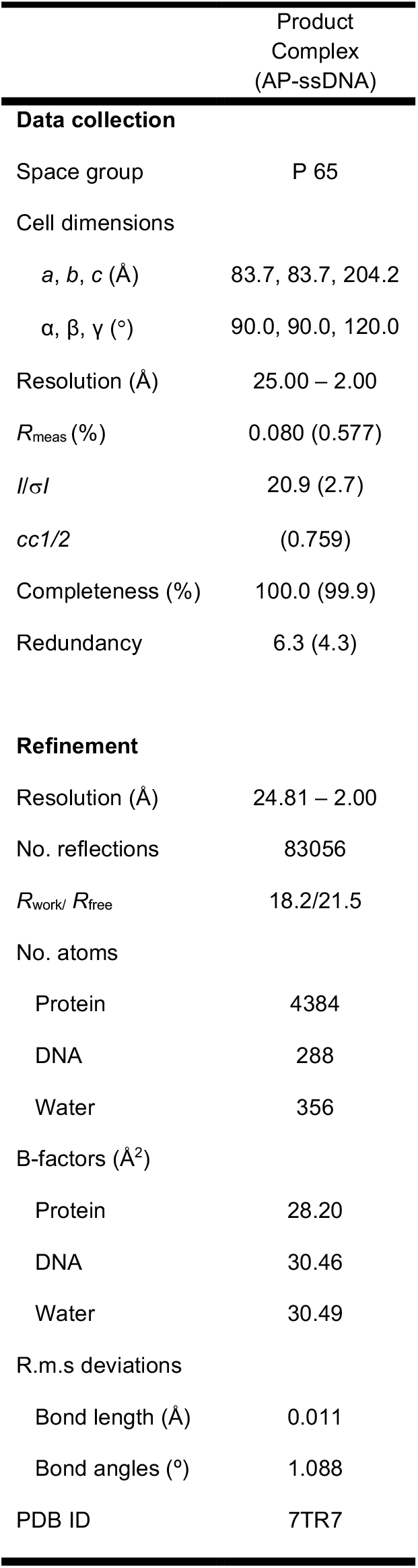
Data collection and refinement statistics. Highest resolution shell is shown in parentheses.

**Figure S3.**
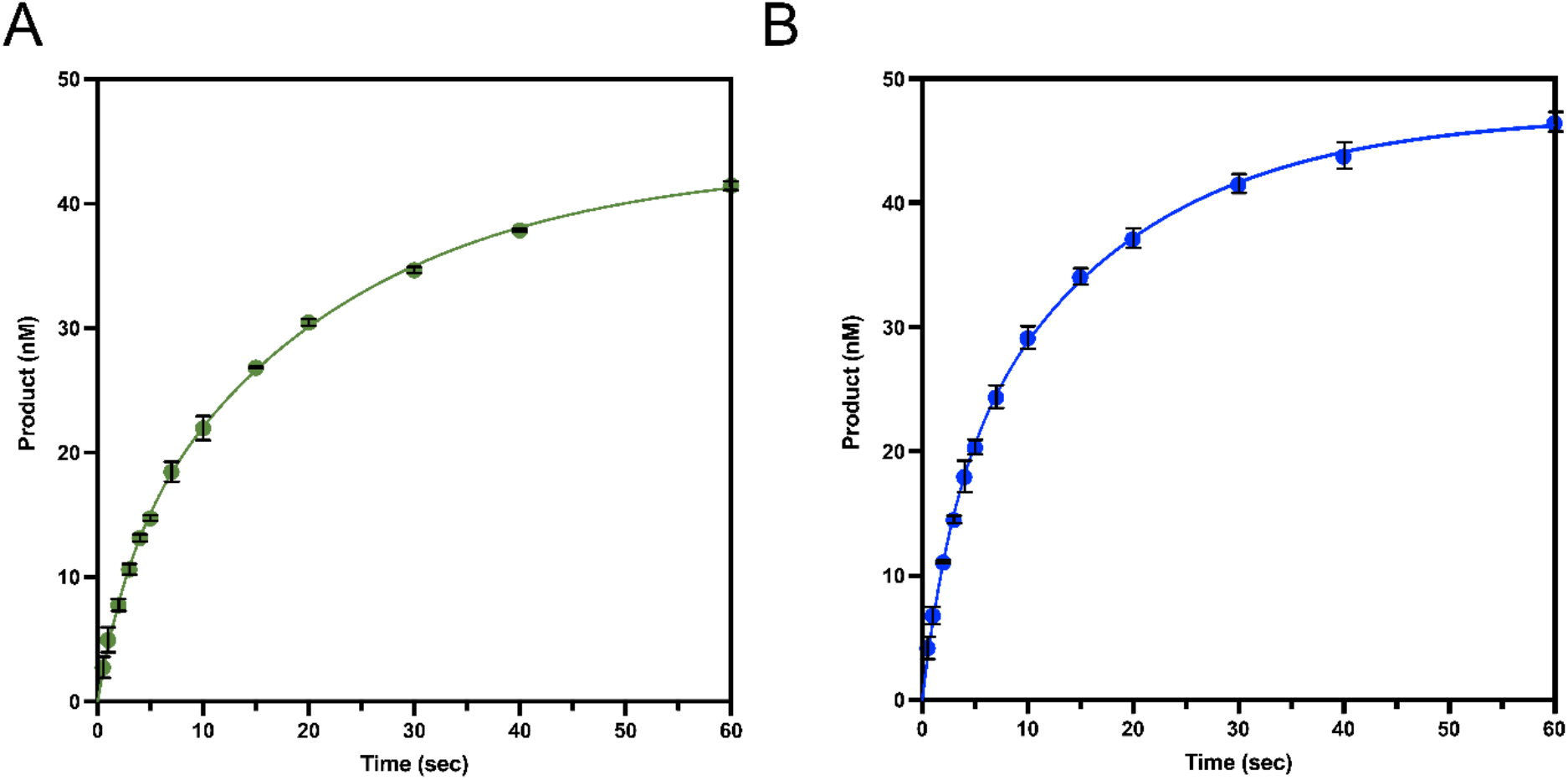
Single turnover kinetics with APE1 R177A. Single turnover kinetic time courses for APE1 R177A with (A) PTJ (green) and (B) Recessed PTJ (blue). All time points are shown as the mean of three independent experiments with error bars (st. dev.) shown. Where error bars are not seen, they are smaller than the data point.

**Figure S4.**
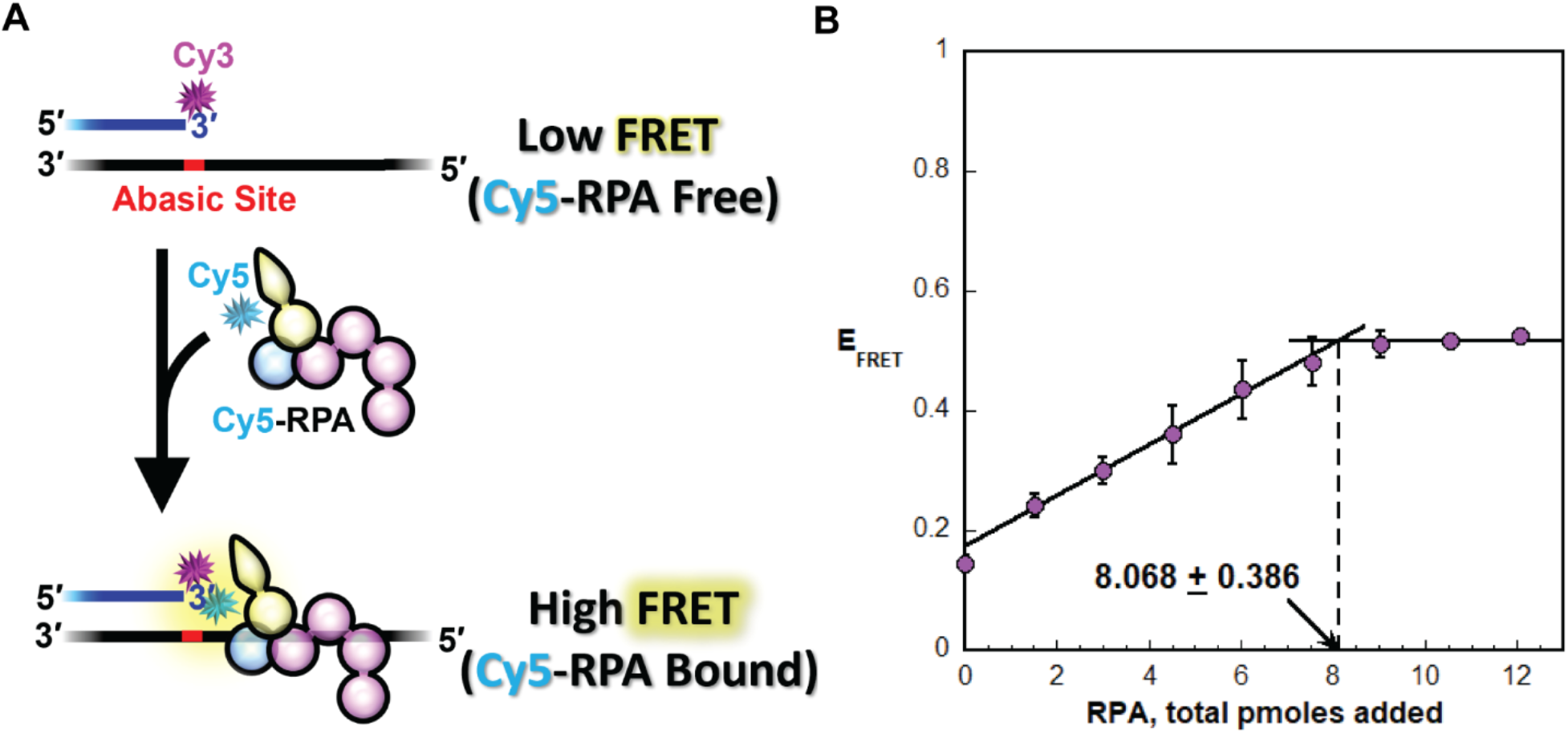
FRET-based titration of Cy5-RPA. The substrate for this assay is a PTJ (**Figure S1**) that contains a Cy3 (FRET donor) fluorophore at the 3′ terminus of the primer strand and an abasic site at the PTJ. (**A**) Schematic representation of experimental procedure. In the absence of Cy5-RPA (FRET acceptor), a FRET is not observed. The ssDNA downstream of the PTJ (33 nt) can accommodate a single Cy5-RPA. RPA engages the ssDNA in an orientation-specific manner such that the Cy5-labeled DBD-D of the RPA32 subunit faces the Cy3 FRET donor on the PTJ, yielding a robust FRET. Hence, addition of Cy5-RPA increases FRET. (**B**) The Cy3-PTJ substrate was titrated with Cy5-RPA and FRET is monitored. The observed E_FRET_ [*I*_665_/(*I*_665_ + *I*_563_)] is plotted as a function of the total amount of added Cy5-RPA and each data point represents the mean ± S.E.M. of at least two independent measurements. Under these experimental conditions, binding is stoichiometric and, hence, FRET increases linearly until the ssDNA is saturated with Cy5-RPA (i.e., equivalence point) (refs). Data is fit to two segment lines (a linear regression with a positive slope and a flat line) and the equivalence point (indicated with standard error of the calculation) is calculated from the intersection of the two segment lines. Saturation is reached at approximately 8 pmole total Cy5-RPA added.

**Figure S5.**
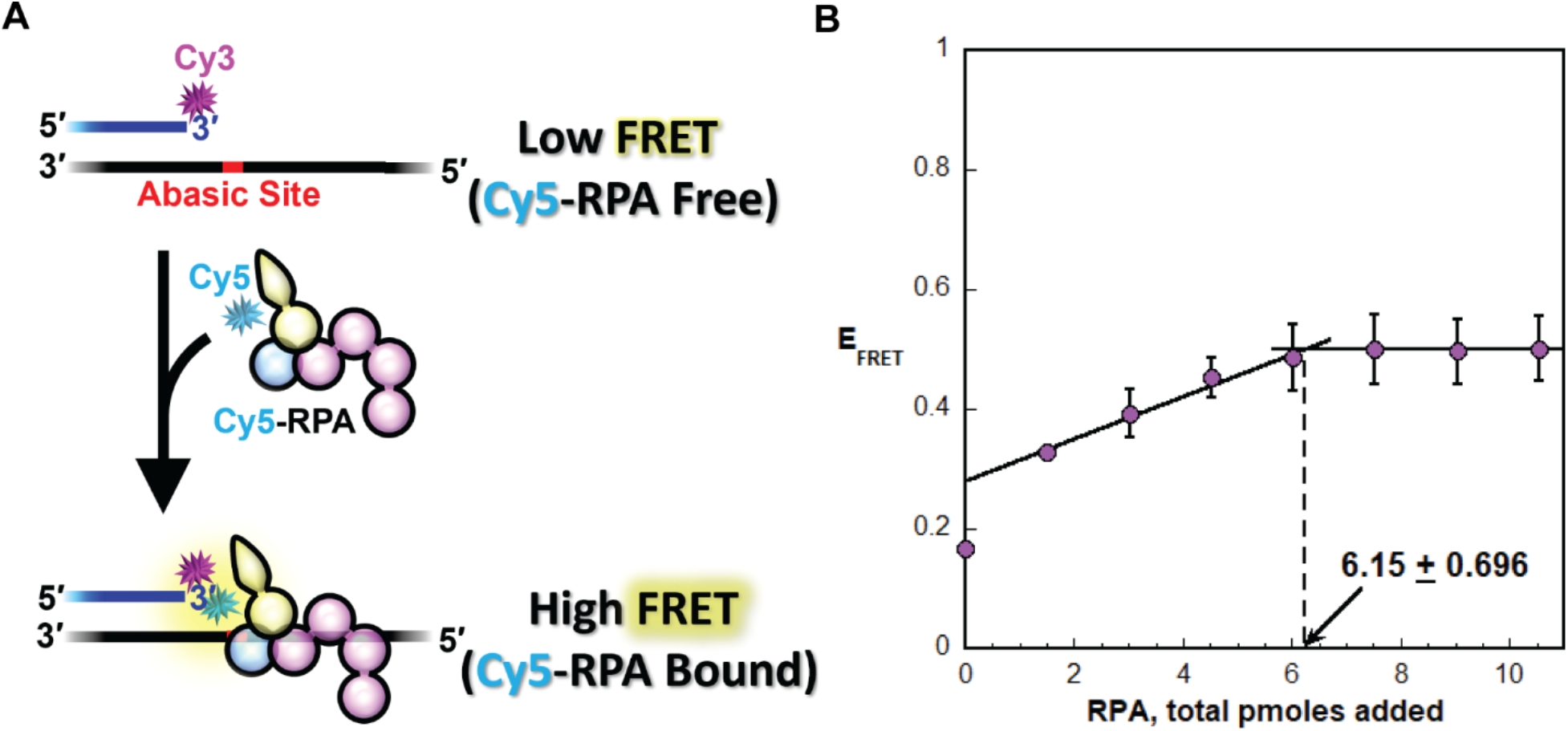
FRET-based titration of Cy5-RPA. The substrate for this assay is a Rec-PTJ (**Figure S1**) that contains a Cy3 (FRET donor) fluorophore at the 3′ terminus of the primer strand and an abasic site 7 nt downstream of the PTJ. (A) Schematic representation of experimental procedure. In the absence of Cy5-RPA (FRET acceptor), a FRET is not observed. The ssDNA downstream of the PTJ (33 nt) can accommodate a single Cy5-RPA. RPA engages the ssDNA in an orientation-specific manner such that the Cy5-labeled DBD-D of the RPA32 subunit faces the Cy3 FRET donor on the PTJ, yielding a robust FRET. Hence, addition of Cy5-RPA increases FRET. (B) Cy3-Rec-PTJ was titrated with Cy5-RPA and FRET is monitored. The observed E_FRET_ [*I*_665_/(*I*_665_ + *I*_563_)] is plotted as a function of the total amount of added Cy5-RPA and each data point represents the mean ± S.E.M. of at least two independent measurements. Under these experimental conditions, binding is stoichiometric and, hence, FRET increases linearly until the ssDNA is saturated with Cy5-RPA (i.e., equivalence point) (refs). Data is fit to two segment lines (a linear regression with a positive slope and a flat line) and the equivalence point (indicated with standard error of the calculation) is calculated from the intersection of the two segment lines. Saturation is reached at approximately 6 pmole total Cy5-RPA added.

**Figure S6.**
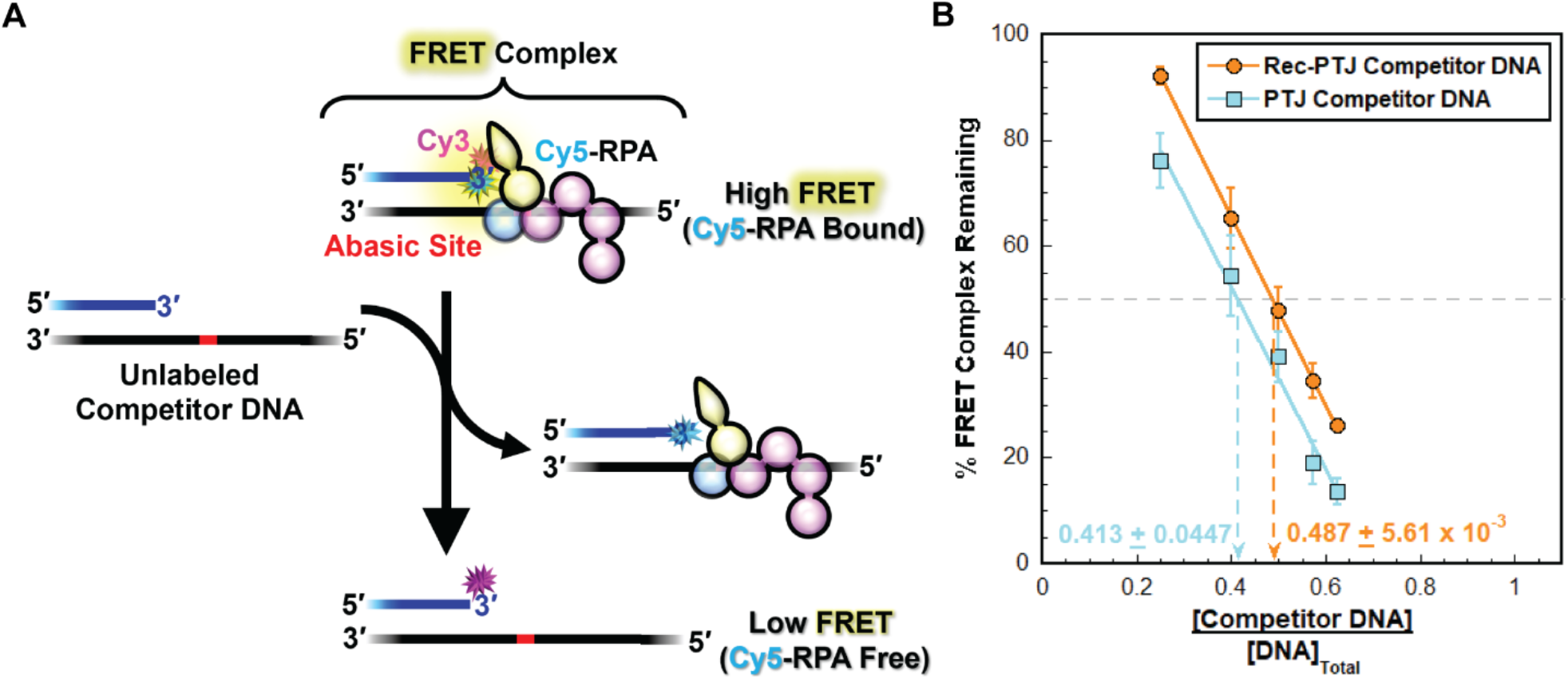
RPA exchange on PTJ DNA containing abasic sites at different positions. (**A**) Schematic representation of the FRET experiment. A 3′ Cy3-labeled Rec-PTJ DNA is pre-saturated with Cy5-labeled RPA and the resultant mixture is then titrated with the respective unlabeled PTJ competitor DNA. E_FRET_ is calculated after each addition of competitor. (**B**) Observed E_FRET_ values for a given competitor DNA are normalized to their respective range and plotted as a function of [Competitor DNA]/[DNA]_Total_ and each data point represents the mean ± S.E.M. of at least three independent measurements. The values observed after the addition of competitor DNA are fit to a linear regression. The [Competitor DNA]/[DNA]_Total_ required for 50% inhibition (i.e., the % FRET Complex remaining decreases to 50%) is calculated from the fit and reported for each competitor DNA.

## Source Data

**Figure 3a – Source Data 1.** Original file for the full and unedited gel of the representative image shown in Figure 3a.

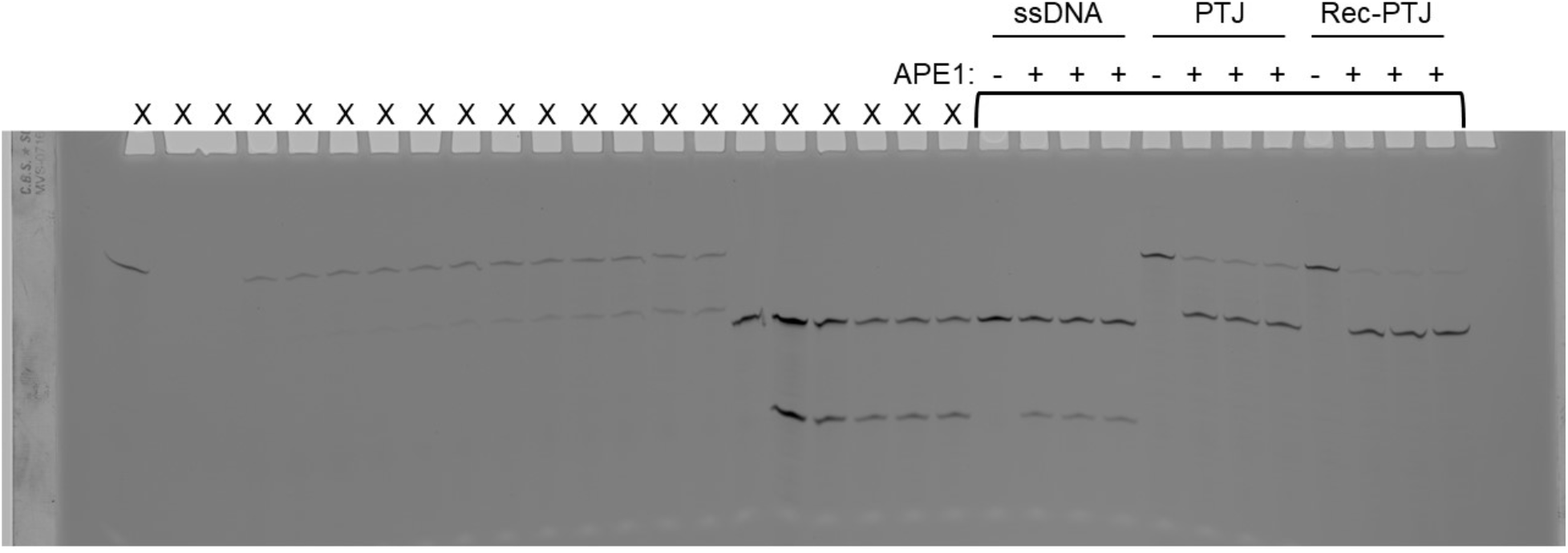

**Figure 3a – Source Data 2.** Uncropped gel of the representative image shown in Figure 3a. Product formation assay showing DNA bands separated on denaturing gel with each DNA substrate in triplicate. Gel lanes labeled with an “X” are not relevant samples.

